# Carbon monoxide-driven bioethanol production operates via a tungsten-dependent catalyst

**DOI:** 10.1101/2024.07.29.605569

**Authors:** Olivier N. Lemaire, Mélissa Belhamri, Anna Shevchenko, Tristan Wagner

## Abstract

Microbial alcohol production from waste gasses is a game changer for sustainable carbon cycling and remediation. While the biotechnological process employing *Clostridium autoethanogenum* to transform syngas (H_2_/CO_2_/CO) is blooming, the reactions involved in ethanol biosynthesis remain to be demonstrated. Here, we experimentally showed that ethanol production initiates via a tungsten-dependent aldehyde:ferredoxin oxidoreductase (AFOR), which reduces acetate to acetaldehyde. Such an unfavourable reaction has often been considered unsuitable for a biological process. To answer this riddle, we demonstrated that the thermodynamic pull of CO-oxidation and ethanol synthesis is crucial for triggering acetate reduction. The experimental setup performed with native CO-dehydrogenase and AFOR highlighted the key role of ferredoxin in stimulating the activity of both metalloenzymes and electron shuttling. The crystal structure of holo AFOR refined to 1.59-Å resolution, together with its biochemical characterisation, provides new insights into the reaction mechanism and the specificities of this enzyme fundamental to sustainable biofuel production.

## Introduction

The transformation of synthetic gases into alcohols is a promising strategy for the carbon cycling economy by converting the greenhouse gas CO_2_ and the polluting carbon monoxide (CO) into added-value molecules. In that regard, *Clostridium autoethanogenum* is already widely used as a bioconverter in industry to convert concentrated waste gasses (H_2_/CO_2_/CO) into acetate, 2,3-butanediol and ethanol (1-8). Despite numerous studies on the topic, there is currently no experimental validation on how CO-dependent ethanol production would operate at the molecular level in *C. autoethanogenum*. Following the most common hypothesis based on genetic and biochemical consideration, the pathway would start with the reduction of acetate by an aldehyde:ferredoxin oxidoreductase (AFOR) and the successive acetaldehyde reduction to ethanol by an alcohol dehydrogenase (2, 5, 9, 10). The first and second reactions would be fuelled by oxidation of the reduced ferredoxin and NAD(P)H pools, respectively. Ferredoxin reduction originates from CO oxidation catalysed by the CO dehydrogenase/acetyl-coenzyme A (CODH/ACS) complex (6), and NAD(P)H is indirectly obtained through ferredoxin oxidation (NADH by the Rnf system, and NAPDH by the Nfn system) (5).

Recent biochemical characterisation of homologous AFOR challenged the plausible role of AFOR in bacterial ethanol production due to the unfavourable reaction of aldehyde generation. As with other acids, acetate reduction requires a very low potential (*E*^0^′=−580 mV for the acetate/acetaldehyde couple), which is difficult to reach for living organisms under standard conditions (11-13). Consequently, the overall reaction would be highly endergonic with regular ferredoxin as an electron donor (considering a redox potential of *E*^0^′=−400 mV (14)). Therefore, it was proposed that AFOR enzymes are physiologically involved in the thermodynamically favourable aldehyde oxidation rather than acid reduction, refuting the previously proposed ethanol production pathways (11-13).

To solve the metabolic riddle of the ethanol production pathway in *C. autoethanogenum*, we characterised the native AFOR by X-ray crystallography, biochemistry and enzymology. On top of providing important information regarding the enzyme activity and mechanism, we showed that the ethanol production pathway is fuelled by CO-oxidation and favoured by the immediate acetaldehyde consumption, sustaining a flux towards alcohol formation and maintaining low intracellular CO concentration.

## Results

### CaAFOR is a W-containing monomeric enzyme inactive as isolated

*C. autoethanogenum* is a strict anaerobe with the stunning chemolithotrophic ability to grow with pure CO as its sole energy and carbon source, suggesting specific metabolic adaptation (6). The bacterium encodes two putative AFOR isoforms, one of them being particularly abundant under CO conditions (15). The bacterium was therefore cultivated on CO to ensure sufficient enzyme quantities for structural and biochemical characterisation. The native anaerobic purification led to a homogenous monomeric population observed by gel filtration and high-resolution native polyacrylamide gel electrophoresis (PAGE, Fig. S1). The protein identity, previously named “AOR1” (8) (WP_013238665, abbreviated as *Ca*AFOR), was confirmed by mass spectrometry (Fig. S2). This protein is similar to the homologues from *Peptoclostridium acidaminophilum* (16) and *Thermoanaerobacter* sp. X514 (13) constituted by a single peptide and proposed to use ferredoxin as an electron donor/acceptor, unlike the heterohexameric NADH-reducing enzymes from *Aromatoleum aromaticum* (11) (Fig. S3 and S4a). These proteins belong to a larger subgroup of proteins also containing the AFOR isoform 1 from *Pyrococcus furiosus*, the historic model in the study of the AFORs and the first to be structurally characterised (17). This group is phylogenetically separated from other types of aldehyde oxidase, such as the formaldehyde:ferredoxin oxidoreductase (FFOR), the benzoyl-CoA reductase BamB or the aliphatic sulfonate:ferredoxin oxidoreductase (ASOR, Fig. S3). All these oxidoreductases contain two metallocofactors, a [4Fe-4S] cluster and a W-containing bis-pyranopterindithiolene cofactor (simplified here as tungstopterin).

The as-isolated *Ca*AFOR did not show any activities when monitoring acetaldehyde oxidation with either methyl viologen (MV) or benzyl viologen (BV), the classic assay used by the community. This phenomenon is known for homologous enzymes (18-20) and was an additional motivation to solve the *Ca*AFOR structure. X-ray fluorescence measurement on protein crystals confirmed the presence of W in *Ca*AFOR (Fig. S5), and the structure was experimentally solved by single-wavelength anomalous diffraction at the W L3 edge. The final model was refined to 1.59-Å resolution (PDB 9G7J, Table S1), the best resolution obtained for this group, granting high confidence to model the metallocofactors (Fig. 1). The asymmetric unit contains two *Ca*AFOR, which have too weak interaction to be stable in solution, hence confirming the monomeric state, unlike structurally characterised homologues (Fig. S4a). However, the overall structure is highly similar to the structures of the homologues, the AFOR from *P. furiosus* being the closest one (Fig. S4b-d). While some positions are perfectly conserved in the active site, which would be candidates to support catalysis, some discrepancies are seen, probably due to fine-tuning of specificities across the different enzymes (Fig. 2a, Fig. S4c and S6a). The active site, embedded in a rather hydrophobic cavity, connects to the solvent by a narrow hydrophobic tunnel, similar to other aldehyde oxidases (Fig. 2b, Fig. S6b). As in homologues, a short distance separates the two cofactors (Fig. 1c and d), allowing efficient electron transfer.

**Fig. 1.**
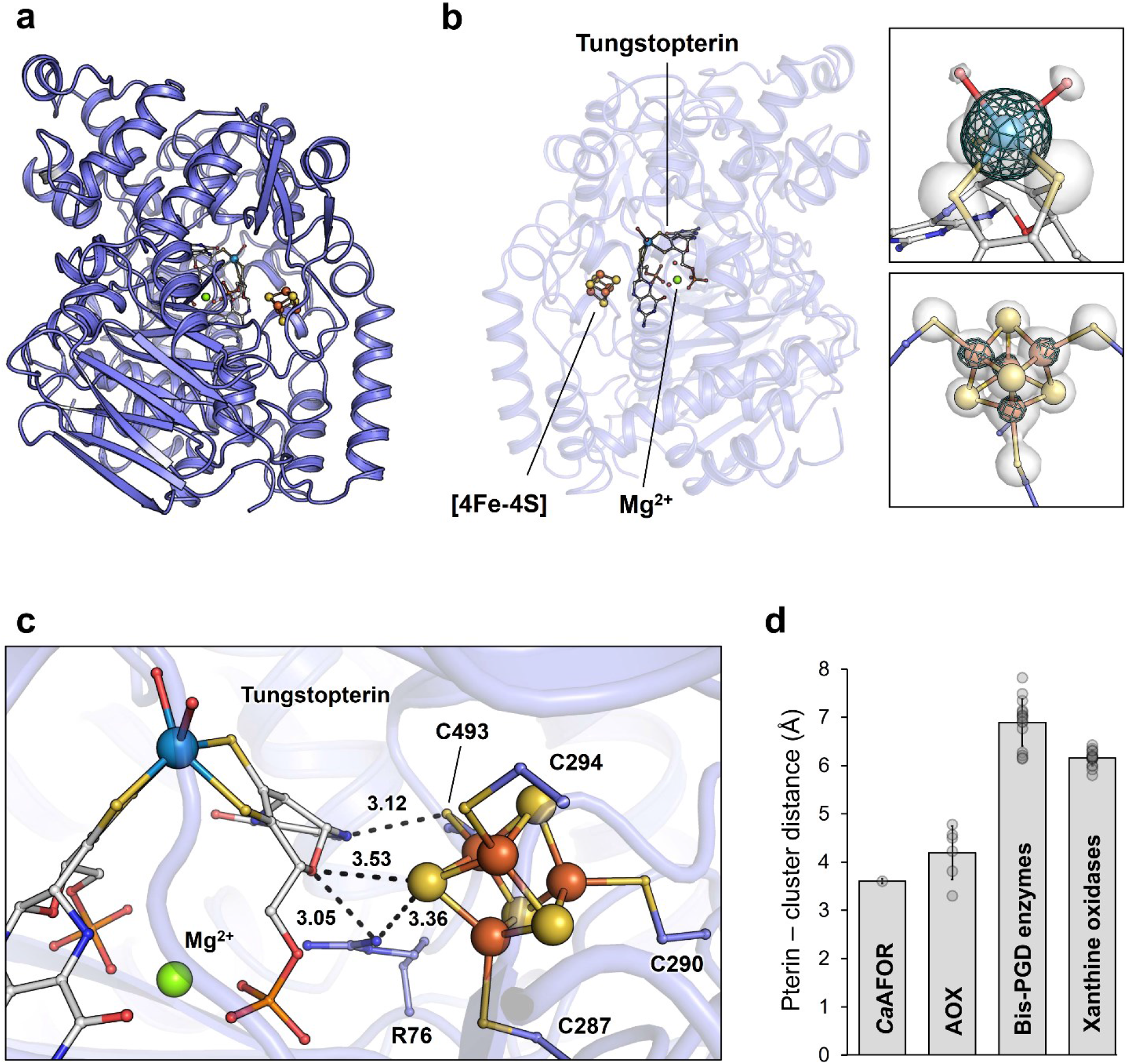
Structure of the AFOR from *Clostridium autoethanogenum*. **a**, Overall structure in a cartoon model. **b**, Metallocofactors composition. Left: *Ca*AFOR is represented as a transparent cartoon model rotated by 180° along the y-axis compared to panel **a**. Ligands are displayed as balls and sticks. Right: The 2*F*_o_-*F*_c_ (1.0 σ) and anomalous map (4.0 σ) around the cofactors are shown as a white transparent surface and a cyan mesh, respectively. **c**, Distances between the [4Fe-4S] cluster and the tungstopterin in *Ca*AFOR are represented in black dash lines and given in Å. **d**, Distances between the [4Fe-4S] cluster and the tungstopterin in structurally characterised pterin-dependent enzymes. AOX stands for all W-dependent aldehyde oxidases (see materials and methods). Average and standard deviation are shown, with individual data shown as transparent grey dots. **a-c**, *Ca*AFOR is shown as a slate blue cartoon with the cofactors and coordinating residues shown as balls and sticks with carbon, oxygen, nitrogen, sulphur, phosphorus, magnesium, iron and tungsten are coloured white (slate for residues), red, blue, light yellow, light orange, green, orange and grey blue, respectively.

**Fig. 2.**
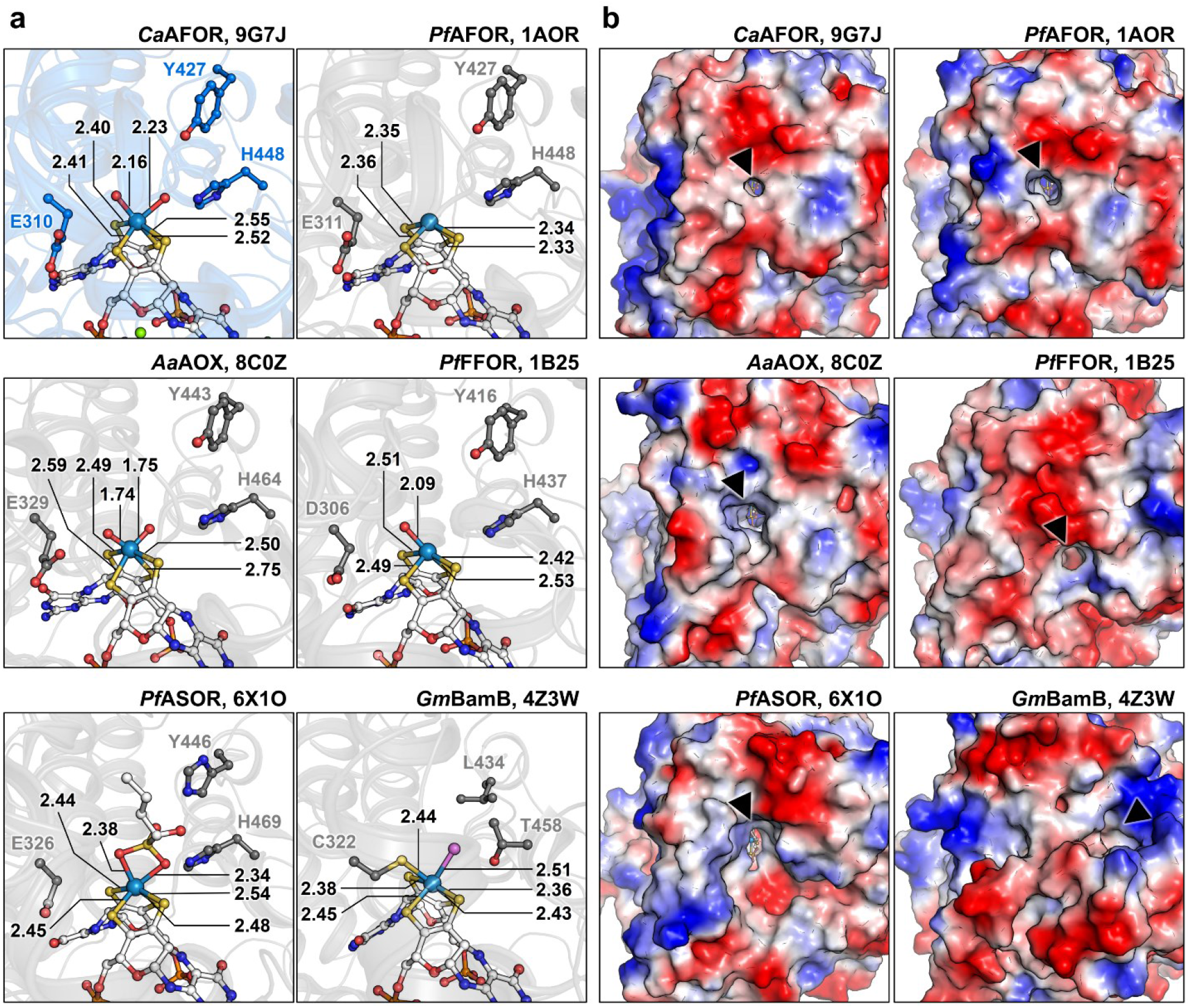
Active site and its entrance across structural homologues. **a**, W-coordination in *Ca*AFOR and structurally characterised homologs. Proteins are shown in transparent cartoons coloured slate (*Ca*AFOR) or grey. Cofactors, coordinating residues and ligands are shown as balls and sticks with carbon, oxygen, nitrogen, sulphur, phosphorus, magnesium, iron and tungsten coloured white (or according to the protein for residues), red, blue, light yellow, light orange, green, orange and grey blue, respectively. The unidentified atom in the BamB structure (4Z3W) is coloured pink. Distance between the W and binding atoms are indicated in Å. **b**, Surface and access to the active site. Proteins are shown as surface coloured by charge distribution from negative (red) to positive (blue). The opening connecting the active site to the solvent is indicated with a black arrow.

The W atom, confirmed by its anomalous signal (Fig. 1b), shows no proteinogenic covalent ligand as all other structurally characterised homologues except the benzoyl-CoA reductase BamB (Fig. 2a). The W atom is coordinated by the dithiolene groups from each pterin and by two atoms modelled as oxygen according to the electronic density (Fig. 1b). A similar ligand has been modelled in the structure of the aldehyde oxidoreductases from *A. aromaticum* (PDB 8C0Z (21)) based on the reanalysed diffraction data from *P. furiosus* enzyme. The W-O distances in the *Ca*AFOR structure are however significantly longer (around 2.2 Å) than the W=O double bond modelled in the structure from *A. aromaticum* (1.75 Å) and closer to the distance of W-OH bonds calculated by QM/MM (2.06 Å (21)) or the W-O-S bonds modelled in the ASOR of *P. furiosus* (PDB 6X1O (22), 2.36 Å, Fig. 2a). Based on the observed distance, the ligands were modelled as hydroxo-rather than oxo-groups (Fig. 1b and 2a). This difference in ligands might, however, be due to an inactivation of *Ca*AFOR that would differ from the active purified enzyme from *A. aromaticum*.

### Ferredoxin is a critical component of *Ca*AFOR activity

The absence of activity of the purified *Ca*AFOR was further investigated. The viologen-based aldehyde oxidase activity, measurable in the cell extract (Fig. S7), exhibited a proportional decrease upon dilution that was not influenced by adding bovine serum albumin (BSA, a molecular crowding agent). The complete loss of viologen-based activity after the first chromatography step and the absence of activity recovery upon chemical treatment with reducing agents (dithionite, Ti(III) citrate), oxidising agent (ferricyanide) or sulphide (Fig. 3a) confirmed the inactivation. This questioned the requirement of an activator present in the cell extract and lost over the first purification step or the suitability of viologen dyes as electron acceptors. Hence, we investigated if the native ferredoxin from *C. autoethanogenum* could be used to monitor aldehyde oxidation instead of artificial dyes.

**Fig. 3.**
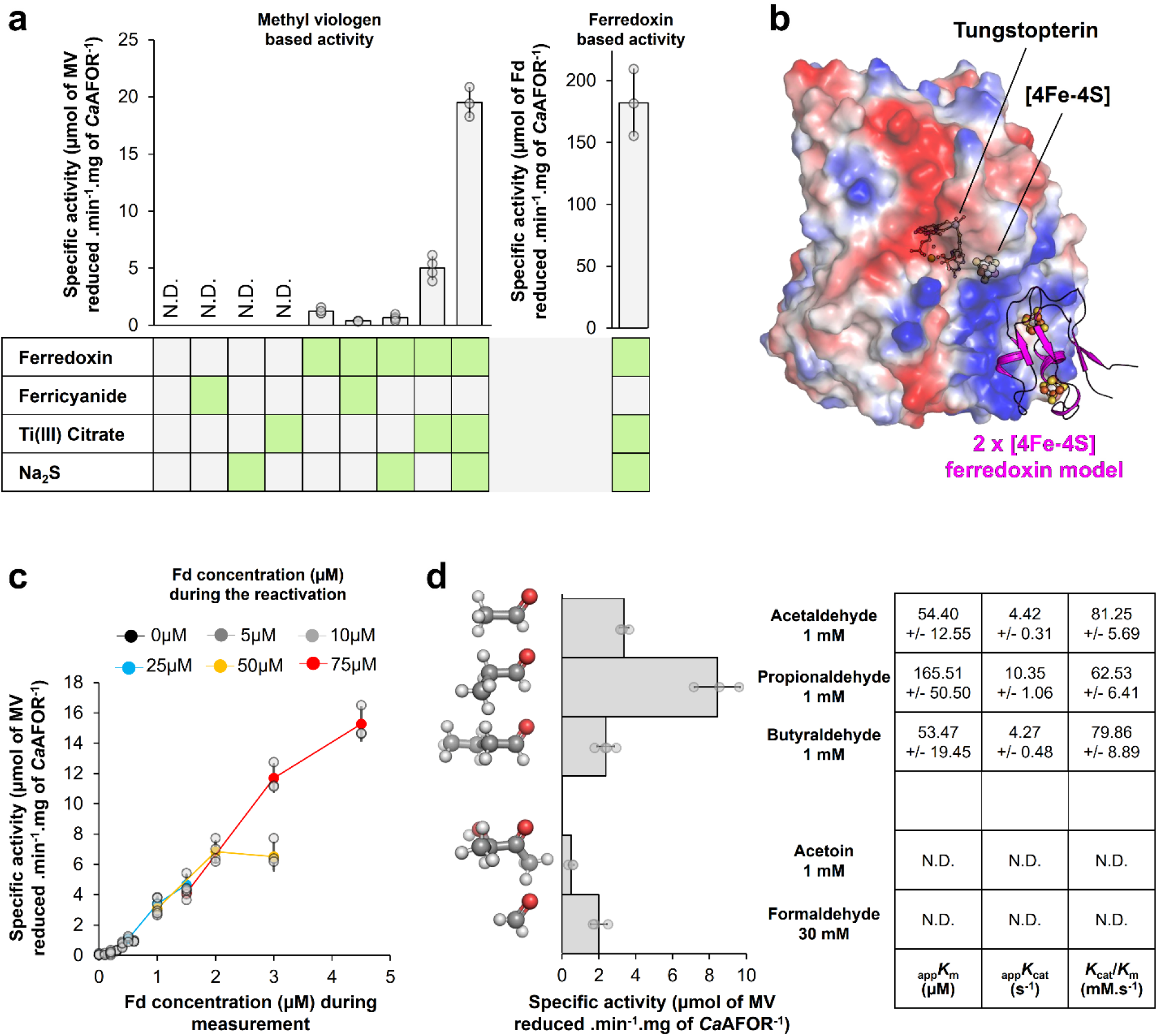
Ferredoxin-dependent AFOR activity. **a**, Left: MV-based *Ca*AFOR activity after incubation with 250 µM ferredoxin, 500 µM ferricyanide, 500 µM Ti(III) citrate and/or 500 µM sulphide. Right: *Ca*AFOR activity with ferredoxin as electron acceptor (after reactivation). Green boxes highlight when the ferredoxin/chemicals are present. **b**, AlphaFold2 model of *Ca*AFOR with the ferredoxin. *Ca*AFOR is displayed as a transparent surface coloured by charge distribution from negative (red) to positive (blue). The ferredoxin is presented as a pink cartoon. Metallocofactors are shown in balls and sticks and coloured as in Fig. 2. **c**, Impact of the ferredoxin concentration on activity measurements. The enzyme was incubated (interpreted here as a “reactivation phase”) with different concentrations of ferredoxin represented by different colours. Three different final concentrations of ferredoxin were assessed during activity measurements for each reactivation procedure. MV serves as electron acceptor. **d**, Aldehyde:MV oxidoreductase activity of the *Ca*AFOR with different aldehydes (shown as balls and sticks with carbon, oxygen and hydrogen coloured as grey, red and white). The kinetic parameters with the different aldehydes are indicated in the table. **a, d**, “N.D.” stands for not detectable. **a, c** and **d**, Average and standard deviation are shown, with individual data shown as transparent grey dots.

With a final purification yield of 614 ± 131 nmol of ferredoxin per gram of soluble proteins obtained after lysis (n=3), the protein is one of the most, if not the most, abundant from *C. autoethanogenum*. As a comparison, the purification yield of CODH/ACS (6), among the most abundant enzymes of the bacterium (6, 15), is around 10-fold lower (55 ± 28 nmol per gram of soluble extract, n=5). Mass spectrometry identification (WP_013236834/OVY50687.1, Fig. S2) did not suggest any other ferredoxin present in the purified preparation. Therefore, the protein is most probably the central physiological electron donor/acceptor of the ferredoxin-dependent reactions occurring in the bacterium and is similar to the characterised homologue from *Clostridium pasteurianum* (WP_003440320, 78.57% identity) used in many studies as an electron acceptor/donor for oxidoreductases (4, 5, 23, 24). An AlphaFold2 prediction of the *Ca*AFOR-ferredoxin interaction displays a docking of the ferredoxin on a positively charged patch near the [4Fe-4S] cluster of the *Ca*AFOR at a distance (11.2-Å) short enough for fast electron transfer (Fig. 3b, Fig. S8a, b and c). Accordingly, the ferredoxin natively isolated in its oxidised state (Fig. S9) was reduced by the purified *Ca*AFOR upon acetaldehyde addition (Fig. S8d).

We noticed that a preincubation phase with ferredoxin led to a detectable MV-dependent acetaldehyde oxidation activity (Fig. 3a). The preincubation effect was enhanced by the presence of a reducing agent such as Ti(III) citrate and, more efficiently, when combined with sulphide (up to 10-fold, Fig. 3a). Using dithionite instead of Ti(III) citrate also enhanced the activity, but the effect was not affected by sulphide addition, probably because the dithionite solution already contained sulphide (Fig. S10a and S11). Replacing MV by BV increased activity by around 2.4 fold, which was expected considering the reaction thermodynamics (*E*^0^′=−374 mV for BV instead of *E*^0^′=−450 mV for MV (25), Fig. S10b). Besides being reproducible, the optimised protocol of preincubation with ferredoxin, Ti(III) citrate and sulphide presented specific activity fluctuations that could not be corrected by BSA (Fig. S10c-d).

Since adding reducing agents strongly stimulated *Ca*AFOR activity during the preincubation phase, we speculated that the as-isolated enzyme was in an inactive oxidised state. To test this possibility, we exposed *Ca*AFOR overnight to air and expected a complete activity loss if the oxidation was irreversible or recovery if the enzyme could still be reduced. O_2_-exposed *Ca*AFOR yielded 45 % of the activity compared to an enzyme stored in an anoxic environment (Fig. S10e), a partial decrease that might be due to the loss of the [4Fe-4S] cluster. While the experiment indicates that enzyme oxidation is reversible, the impact of the preincubation phase on the enzyme is more complex to interpret and might hide a more intricate reactivation process in which ferredoxin association plays a key role.

To decipher the effect of ferredoxin on *Ca*AFOR activity, we measured the ferredoxin reduction AFOR rates instead of using viologen as an electron acceptor. After enzyme reactivation, we measured a specific activity of 182.12 ± 26.97 µmol of reduced ferredoxin per min per mg of *Ca*AFOR (Fig. 3a), a high activity for this type of enzyme (11, 13, 16, 19, 26-30). The 10-fold difference observed between the reduction rates of MV and ferredoxin contradicts the possibility that ferredoxin only serves as an electron bridge between AFOR and MV. It rather strengthens the hypothesis of a role that might indeed contribute to *Ca*AFOR reactivation by stimulating the reduction of tungstopterin. To deepen our investigation, different concentrations of ferredoxin were tested during both reactivation and activity measurements. Up to 75 µM, the activity increases linearly with the concentration of ferredoxin used during reactivation (Fig. 3c). Higher ferredoxin concentrations did not enhance enzyme reactivation (Fig. S10f). The ferredoxin concentration during activity measurement also positively influences the measured activity, this later increasing with the ferredoxin concentration up to 5-10 µM ferredoxin during activity measurement. A large excess of ferredoxin is hence necessary for a maximal reactivation of the enzyme, but a relatively high ferredoxin concentration is also important to maintain *Ca*AFOR activity. This echoes with the large excess of ferredoxin necessary for maximal activity of the enzyme of *Thermoanaerobacter* sp. X514 (13) and explains the loss of *Ca*AFOR activity in the extract upon dilution, as it decreases the concentration of native ferredoxin.

### *Ca*AFOR substrate specificity

The affinity of *Ca*AFOR for different aldehyde was tested. Because of the observed fluctuations during the reactivation procedure (Fig. S10c) and the known unstable properties of aldehydes, the estimated apparent kinetic parameters for each substrate exhibited relatively large deviations. The results indicated that *Ca*AFOR catalyses the oxidation of acetaldehyde, propionaldehyde and butyraldehyde with a turnover and affinity in the same order of magnitude (Fig. 3d). The enzyme turnover and _app_*K*_m_ with these substrates is in the range that is determined by characterised homologues (11, 13, 19, 20, 28, 30). This relative non-specificity is a common feature of the enzyme family (13, 16, 26, 30). Still, *Ca*AFOR poorly catalysed formaldehyde oxidation, requiring concentrations above confident measurements to extract kinetic parameters. Using branched aldehyde such as acetoin as substrate only yields an extremely low activity, which might be due to the characteristic of the tunnel specificities (Fig. 2b and Fig. S6b). The hydrophobicity of the tunnel could limit the efficient diffusion of the small, relatively polar formaldehyde, while its narrow diameter may prevent that of branched aldehydes.

The homologue from *A. aromaticum* (29) has recently been shown to exhibit hydrogenase activity. Here, we could not observe H_2_ oxidation with either MV or ferredoxin as an electron acceptor in our conditions for *Ca*AFOR. Accordingly, H_2_ oxidation in *C. autoethanogenum* is operated by dedicated hydrogenases that have been suggested to catalyse the reverse reaction during the CO-dependent growth (4). Hence, our data do not support any hydrogenase activity for the *Ca*AFOR or any direct function in H_2_ metabolism.

### CO-driven alcohol production

Following our hypothesis, *Ca*AFOR should be the starting point of the ethanol pathway by generating acetaldehyde from acetate. As expected, no reliable *in vitro* acetate reduction by *Ca*AFOR could be monitored with either reduced MV, BV or ferredoxin due to the unfavourable thermodynamic of the reaction (Fig. 4a). The reaction equilibrium is probably reached before a reliable activity could be monitored. *In vivo*, the physiological context is drastically different as the favourable acetaldehyde reduction to ethanol coupled to NAD(P)H oxidation would prevent the aldehyde accumulation while the ferredoxin pool would be maintained in a reduced state by CO oxidation (Fig. 4a and b). Therefore, the overall process from acetate to ethanol would be largely exergonic. To mimic these metabolic pulls, *Ca*AFOR was coupled to the NADH-dependent alcohol dehydrogenase (ADH) from *Saccharomyces cerevisiae* and the native CODH/ACS complex (6) to fuel the acetate-reducing reaction with electrons from CO-oxidation (Fig. 4b). The experiment was performed sequentially by testing the compatibility of the three different functional modules (electron donating, acetate reduction, and acetaldehyde elimination).

**Fig. 4.**
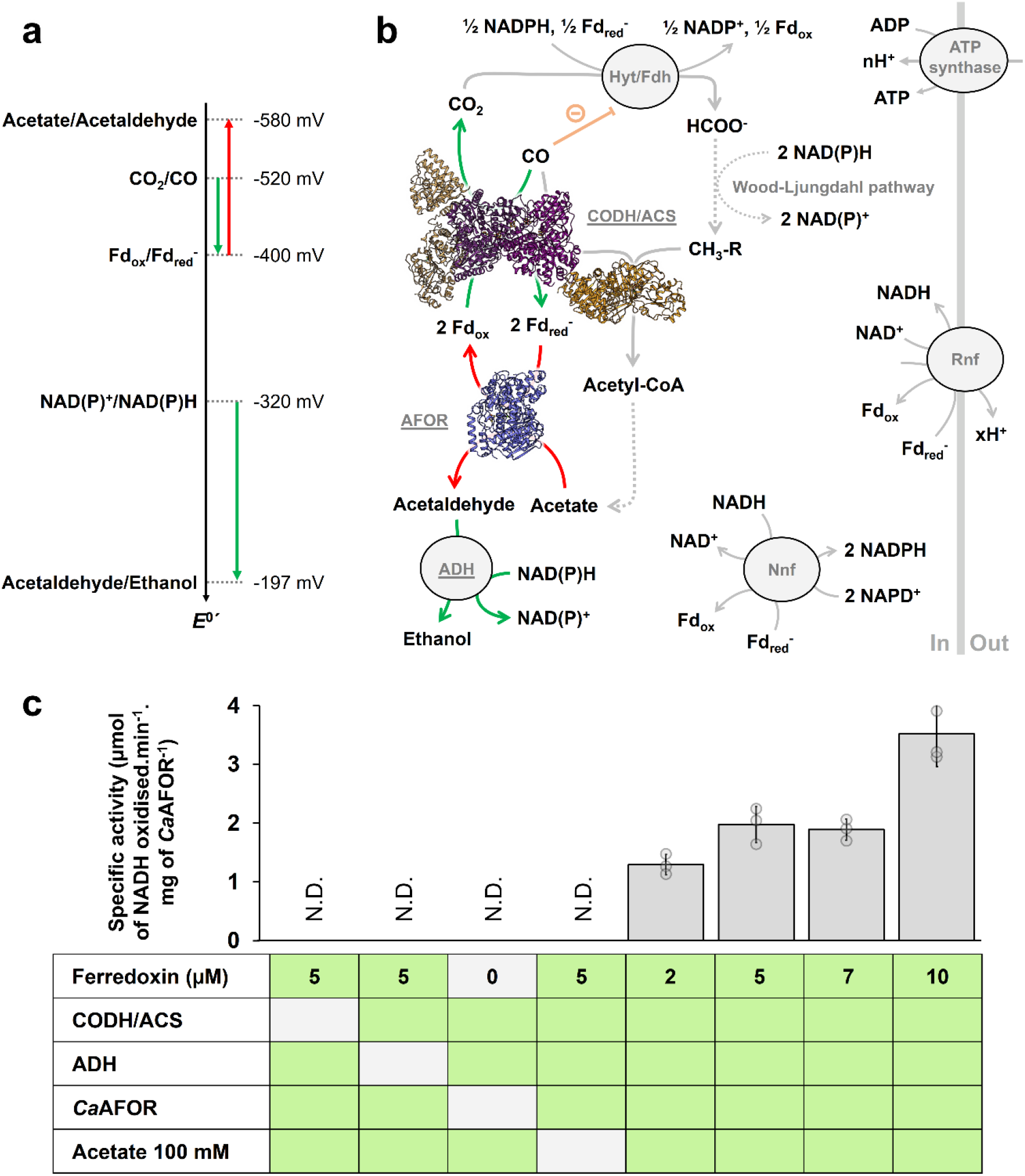
Coupled enzymatic assay for *in vitro* CO-dependent ethanol production. **a**, Standard redox potential of the couples involved in ethanol production. Thermodynamically favourable and unfavourable reactions are shown by green and red arrows, respectively. **b**, Metabolic pathway for ethanol production in *C. autoethanogenum*. The experimental CODH/ACS and *Ca*AFOR structures are shown in cartoon (6). CO inhibition affecting the electron confurcating Hyt/Fdh complex is coloured orange. Dotted lines indicate multi-step reactions. Enzymatic reactions used in the coupled assay are coloured green and red, as for panel **a** (ADH from *S. cerevisiae* is being used instead of *Ca*ADH). The number of proton translocated through ferredoxin oxidation and for ATP synthesis were not experimentally determined and therefore are noted x and n, respectively. **c**, Specific activity in the different conditions for the coupled assay. Average and standard deviation are shown, with individual data shown as transparent grey dots. “N.D.” stands for not detectable. Green boxes contain the proteins/acetate.

We previously noticed that the CODH/ACS complex from *C. autoethanogenum* loses drastic activity during purification (6). When the reactivation procedure developed for the *Ca*AFOR was applied to the purified CODH, a 150-fold increase in MV-based CO-oxidation activity was measured (Fig. S12a). Loss of activity over time was, however, still observable despite the enzyme reactivation. Unlike the *Ca*AFOR, the reactivation of the CODH activity is sulphide-independent, and the activity is relatively similar if MV or ferredoxin is employed as the final electron acceptor (Fig. S12b and c).

The CO-driven ethanol synthesis was performed with CODH/ACS and ADH in saturating concentration, such as the *Ca*AFOR would be the rate-limiting enzyme in the process. Firstly, we confirmed that AFOR and acetaldehyde:NAD^+^ oxidoreductase activities were not impacted by the addition of CO (104.6 ± 10.0 and 92.5 ± 18.0 % of the activity with around 33 % CO in the gas phase, respectively). Secondly, we confirmed the absence of a NADH oxidase background when all components were added except acetate. The oxidation of NADH reached a robust, specific activity when all components were added to the coupled assay, indicating that the *Ca*AFOR produces acetaldehyde under this condition (Fig. 4c). The NADH oxidation rate was dependent on CO concentration, as around 3 % final CO in the gas phase was not sufficient to detect reliable activity, and was impacted by ferredoxin concentration (Fig. 4c). It cannot be deciphered if the effect of ferredoxin concentration is due to *Ca*AFOR or CODH reactivation, electron transfer from CODH/ACS to *Ca*AFOR, or both. The kinetic parameters for acetate were affected by the pH, with the enzyme exhibiting higher rates at acidic pH (Fig. S13). The maximal monitored acetate reduction turnover by *Ca*AFOR was 9.20 ± 0.74 s^-1^, comparable with rates obtained during the thermodynamically favourable aldehyde oxidation with viologen derivatives catalysed by *Ca*AFOR or homologues (13, 16, 26), and around 1,000 fold higher than the acid reduction turnover determined with the closely related enzyme from *Thermoanaerobacter* sp. X514 (13).

## Discussion

The carbon cycling economy is one of the most important challenges that our modern society faces to counterbalance greenhouse gas emissions and ensure long-term sustainability. An existing solution is to rely on the chemistry of acetogenic bacteria to convert industrial greenhouse waste gases (e.g., syngas released from the steel mill) into alcohols used as biofuels or as building blocks for organic chemistry. While *C. autoethanogenum* is in the spotlight of this technology, its highly specialised CO-metabolism remains to be unveiled at the molecular level. While we previously unravelled how the CO-capturing strategy of the CODH/ACS complex operates via a porous network (6), we describe in this work the acetate reduction step catalysed by a W-dependent AFOR for ethanol production.

By performing the reversible conversion of different aldehydes into acids, AFORs are environmentally and biotechnologically important, with a recent application in synthetic biology (29, 31). Numerous mechanisms have been proposed based on structural and biophysical characterisation or on molecular simulations, (21, 22, 32, 33). In all of them, the conserved glutamate and histidine (Fig. 2a, Fig. S4c and S6a) would assist the reaction for proton exchange, while the reaction intermediate and the reported hydroxyl/oxo ligands on the tungsten together with its redox state differ. The near-atomic resolution structure of the *Ca*AFOR provides additional highlights that would challenge a reaction involving two oxo axial ligands. First, the distance of the catalytic W atom and both axial ligands better fit with hydroxyl groups in the as-isolated *Ca*AFOR structure. Then, one of the hydroxyl groups establishes short hydrogen bonds with the perfectly conserved glutamate (Glu312) and histidine (His448, Fig. S6a), a position that should be occupied by the acetaldehyde/acetate molecules and reaction intermediates. Finally, since the enzyme requires reducing conditions to restore its full catalytic potential, it is expected that upon successive reduction the W(IV) would potentially lose one of the axial ligands and would have a vacant position to bind the substrate. However, as the structure reflects an inactive state, it is also possible that upon reactivation slight rearrangements of the active site and axial ligand would occur to stabilise a state previously described in the literature.

Compared to other oxidoreductases that can be reactivated with only reducing agents (e.g., CODH (34) or hydrogenase (35)), *Ca*CODH and *Ca*AFOR additionally require ferredoxin. We suspect that ferredoxin docking would provide an electron path from Ti(III) citrate to the metallocenter via the [4Fe-4S]-cluster without provoking large conformational rearrangements due to the robustness of the protein scaffold. One key difference between the two enzymes is the stimulation of sulphide in the case of *Ca*AFOR. A ferredoxin-independent reactivation by sulphide was previously described for the FFOR from *P. furiosus* (18), and dithionite (a sulphide source) is regularly used in the purification of AORs to preserve the activity (18, 19). The genomic environment of the gene coding for *Ca*AFOR contains a gene coding for a putative molybdo/tungstopterin cofactor sulfurase, homologous to the gene responsible for sulphur addition on the Mo atom in xanthine oxidases (36) (Fig. S14a). Hence, in its physiological state, this AFOR might present at least one sulphur-based ligand (sulfhydryl/sulfido) on the tungsten, similar to the protein belonging to the DMSO reductase family (e.g., formate dehydrogenases (37)) and Mo-dependent aldehyde oxidase (38). The progressive oxidation of the ligand during the purification would explain the gradual inactivation previously observed (17-19, 39) and the presence of oxygen rather than sulphur in the presented structure (Fig. S14b). Despite our efforts to soak or co-crystallise the enzyme with ferredoxin, dithionite and/or sulphide, no significant modification of the electronic density could be observable in the crystal structure, an effect most probably due to the crystal packing preventing interaction with the ferredoxin *in cristallo*. Further investigation by complementary approaches such as spectroscopy will have to investigate if a sulphur atom has a role in the reactivation or reaction mechanism. Obtaining additional data on the reactivated enzyme will also shine light on the pre-catalytic state, which might corroborate the recent work performed on the enzyme from *A. aromaticum* or present an unexpected reconfiguration of the W environment.

The AFORs has been shown to artificially reduce acids *in vitro*, but due to the low measured rates (29, 31) and the poor affinity of the characterised enzymes for acids, this reaction has sometimes been claimed as physiologically irrelevant (11-13). Nevertheless, when mimicking the metabolic context by constantly overflowing the system with CO-dependent ferredoxin reduction and eliminating the product with an ADH, we showed that *Ca*AFOR catalyses a robust aldehyde production (Fig. 4c), strengthening its role in alcohol production as previously suggested in acetogenic bacteria and as demonstrated in archaea (8-10). The high _app_*K*_m_ of *Ca*AFOR for acetate and the acidic pH dependency can be rationalised as follows: (i) at lower pH the redox midpoint potential of the acetate/acetaldehyde couple will increase, making acetate reduction thermodynamically more favourable; (ii) the stimulation of the measured rates at pH 5.5 and the hydrophobic nature of the cleft and active site would indicate that the enzyme prefers acetic acid as substrate rather than acetate, as previously proposed (5, 26); (iii) the intracellular pH of the bacterium has been estimated to be around 6.0 (5), also facilitating acetate protonation; (iv) the bacterial growth can tolerate concentration of acetate reaching the hundreds of millimolar range (2, 3, 8, 40), and it was previously proposed that internal acetate concentration might be higher than that detected in the culture medium (5). The ethanol production, requiring reducing equivalent, could also be triggered by the accumulation of acetate to prevent jamming of the energetic metabolism and/or acidification of the medium (41). In addition, *Ca*AFOR will act as an electron exhauster, generating oxidised ferredoxin to maintain high CO oxidation rates and avoiding the accumulation of the toxic molecule in the cell that would poison acetogenesis by direct inhibition of one of its initial steps (4) (Fig. 4b). This is coherent with the CO tolerance of the *Ca*AFOR and the high CO concentration required for the coupled enzyme assay. With this dual function of controlling the reduced ferredoxin pool and eliminating poisonous high acetate concentration, *Ca*AFOR could represent a key adaptation to CO-metabolism while providing a fantastic opportunity for bioethanol production.

## Methods

### Bacterial strains and growth conditions

*C. autoethanogenum* DSM 10061 was obtained from the Deutsche Sammlung von Mikroorganismen und Zellkulturen GmbH, Braunschweig, Germany. *C. autoethanogenum* was cultivated under strict anaerobic conditions at 37 °C in CaGM growth medium (2) in which Ti(III) citrate was omitted, as previously described (6). Cells were grown chemolithotrophically under agitation (140 rpm) in 1 l Duran bottles containing 0.5 l medium (pH 6.0) and a gas phase containing 100 kPa of 100% CO. Cell growth was monitored spectrophotometrically by measuring the optical density at 600 nm (OD600), and changes in headspace pressure. The cells were harvested in the late-exponential phase by centrifugation and kept frozen at −80 °C in anaerobic conditions under 100 kPa of H_2_/CO_2_ (80:20%). For purification of CODH/ACS and ferredoxin, heterotrophically-gown cells were used. In these cultures, *C. autoethanogenum* was grown in a 10 l fermenter in 9 l of a modified CaGM medium containing only 2 g.l^-1^ 2-(N-morpholino)ethanesulfonic acid (MES) buffer and 5 g.l^-1^ fructose, operating at 37°C and 150 rpm. The fermenter was inoculated with 0.5 l of a heterotrophic culture of *C. autoethanogenum* (OD_600_ around 2.5) grown with 5 g.l^-l^ fructose and a gas-phase containing 100 kPa of H_2_/CO_2_ (80:20%). After 24 h of culture under continuous bubbling with 100% N_2_, 50 ml of an anoxic solution of cysteine/HCl 4% (w/v) was added, and the bubbling gas was switched to H_2_/CO_2_ (80:20%). Cells were harvested after 8 h. A fermenter usually reached an OD_600_ of 2 and yielded 60 g of cells (wet weight).

### Protein purification

Cell lysis and preparation of extracts were performed in an anaerobic chamber filled with an N_2_/CO_2_ atmosphere (90:10%) at room temperature. The cells were lysed *via* three rounds of French Press at around 1000 PSI (6.895 MPa). To guarantee minimum oxygen contamination, the French press cell was prior flushed with N_2_ and washed twice with anoxic buffer. Soluble extracts were prepared by ultracentrifugation at 185,500 x *g* for 1 h at 4 °C. Protein purification was carried out under anaerobic conditions in a Coy tent with a N_2_/H_2_ atmosphere (97:3%) at 20 °C and under yellow light. Samples were passed through a 0.2 μm filter prior to loading on chromatography columns. During purification, multi-wavelength absorbance monitoring (at 280, 415 and 550 nm) and sodium dodecyl sulphate polyacrylamide gel electrophoresis (SDS-PAGE) were used to follow the enzymes.

The purification of the native *Ca*AFOR and ferredoxin was originally performed based on their colour, their expected size on SDS PAGE and the natural abundance of the proteins. They were purified several times at 20°C under strict exclusion of oxygen and under yellow light with a similar, reproducible protocol. About 15 g (wet weight) of frozen cells were thawed and diluted with 15 ml of 50 mM Tris/HCl pH 8.0 and 2 mM dithiothreitol (DTT) before lysis. After lysis, the soluble extract was diluted with 125 ml of the same buffer. The sample was loaded on 3 × 5 mL HiTrap™ DEAE Sepharose FF (GE Healthcare, Munich, Germany) equilibrated with the same buffer. After a 5-column volume (CV) washing, proteins were eluted with a 0 to 0.4 M NaCl linear gradient for 13 CV at a 2 ml.min^-1^ flow rate. The *Ca*AFOR eluted between 0.16 M and 0.19 M NaCl. Two volumes of 50 mM Tris/HCl pH 8.0 and 2 mM DTT were added to the pooled fractions before loading on a 5 ml HiTrap™ Q-Sepharose High-Performance column (GE Healthcare, Munich, Germany) column equilibrated with 25 mM Tris/HCl pH 7.6 and 2 mM DTT. After a 5 CV washing with the same buffer, proteins were eluted with a 0.05 to 0.45 M NaCl linear gradient for 12 CV at a 1 ml.min^-1^ flow rate, the protein of interest eluting between 0.19 and 0.24 M NaCl. The resulting pooled fraction was diluted with one volume of 25 mM Tris/HCl pH 7.6, 2 M (NH_4_)_2_SO_4_, and 2 mM DTT and loaded on a 5 ml HiTrap™ Phenyl Sepharose High-Performance column (GE Healthcare, Munich, Germany). The sample was eluted with a 1.0 to 0.4 M linear gradient of (NH_4_)_2_SO_4_ for 8 CV at a flow rate of 1 ml.min^-1^. The protein was eluted between 0.73 and 0.48 M (NH_4_)_2_SO_4_. The resulting pooled fraction was diluted with two volumes of 25 mM Tris/HCl pH 7.6, 2 M (NH_4_)_2_SO_4_, and 2 mM DTT and loaded on a Source™ 15PHE 4.6/100 PE (GE Healthcare). The sample was eluted with a 1.75 to 0.8 M linear gradient of (NH_4_)_2_SO_4_ for 23 CV at a flow rate of 1 ml.min^-1^. The protein was eluted between 1.73 and 1.56 M (NH_4_)_2_SO_4_. After fraction concentration on 10-kDa cut-off centrifugal concentrator (nitrocellulose, Vivaspin from Sartorius), contaminants were separated by size exclusion chromatography on a Superdex 200 Increase 10/300 GL (GE Healthcare, Munich, Germany) in 25 mM Tris/HCl pH 7.6, 10% (v/v) glycerol and 2 mM DTT with a flow rate of 0.4 ml.min^-1^. The *Ca*AFOR was eluted with a 13.40 ml elution volume. The pooled fraction was concentrated, and the protein was directly used for crystallisation or stored at −80 °C under anoxic conditions.

The ferredoxin eluted between 0.27 M and 0.33 M NaCl at the DEAE chromatography step. Two volumes of 50 mM Tris/HCl pH 8.0 and 2 mM DTT were added to the pooled fractions before loading on a 5 ml HiTrap™ Q-Sepharose High-Performance column (GE Healthcare, Munich, Germany), equilibrated with 25 mM Tris/HCl pH 7.6 and 2mM DTT. After a 3 CV washing with the same buffer, proteins were eluted with a 0.2 to 0.5 M NaCl linear gradient for 8 CV at a 1 ml.min^-1^ flow rate, the protein of interest eluting between 0.36 and 0.40 M NaCl. 2.5 M (NH_4_)_2_SO_4_ was added as anoxic powder to the resulting pooled fraction, and the sample was loaded on a 5 ml HiTrap™ Phenyl Sepharose High-Performance column (GE Healthcare, Munich, Germany). The sample was eluted at 1 M (NH_4_)_2_SO_4_ at a flow rate of 1 ml.min^-1^. The protein was concentrated on a 3-kDa cut-off centrifugal concentrator (nitrocellulose, Vivaspin from Sartorius), and the buffer was exchanged for 25 mM Tris/HCl pH 7.6, 10% (v/v) glycerol and 2 mM DTT. The protein concentration could not be estimated using the Bradford method and was estimated by the absorbance of 390 nm of the aerobically oxidised protein using a molar extinction coefficient of 30,000 M^-1^. cm^-1^ (42). The protein was directly used or stored at -80 °C under anoxic conditions.

The CODH/ACS purification protocol was previously published (6).

### Mass Spectrometric Analysis

Proteins were in-gel digested with trypsin (Promega, Germany). The resulting peptide mixtures were analysed by LC-MS/MS on a RSLCnano system UltiMate^®^ 3000 serie interfaced on-line to an Orbitrap HF hybrid mass spectrometer; the nano-LC system was equipped with Acclam PepMaptm 100 75 µm x 2 cm trapping column and 50 cm μPAC analytical column (all Thermo Fischer Scientific, Bremen, Germany). Peptides were separated using 75 min linear gradient, solvent A -0.1 % aqueous formic acid, solvent B -0.1 % formic acid in acetonitrile. Data were acquired in DDA mode using Top20 method, precursor m/z range was 350-1600; resolution – 120000 and 15000 for precursor and fragments respectively; dynamic exclusion time was set on 15 s. Acquired spectra were matched by Mascot software (v.2.2.04, Matrix Science, UK) with 5 ppm and 0.025 Da tolerance for precursors and fragments against *C. autoethanogenum* protein sequences in NCBI database (August 2024, 23,655 entries), the results were evaluated by Scaffold software (v.5.3.3, Proteome Software, US) using 99 % and 95 % protein and peptide probability thresholds, and FDR (False Discovery Rate, Scaffold option) calculated <1 %.

### Sulphide quantification

The sulphide concentration in the dithionite solution was quantified by a protocol adapted from the diamine method (43). Shortly, different volumes of a freshly prepared anoxic 5 mM sodium dithionite solution (in 50 mM Tris/HCl pH 8.0) were sampled and dissolved in aerobic 1 % (w/v) zinc acetate solution in glass tubes (final volume 1 ml) to form zinc-sulphide precipitates. Titration of sulphide was performed by the addition of 80 µl of the diamine reagent (4 g/l *N,N*-dimethyl-1,4-phenylenediamine sulphate, 6 g/l FeCl_2_·H_2_O, dissolved in a cold 12 M HCl solution). The reaction was performed for 30 minutes in the dark before the spectrophotometric measurement of the reaction product at 670 nm. Sulphide concentration was determined by comparing it with the standard of an anoxic pure sulphide solution.

### Phylogeny analysis

The protein sequence of *Ca*AFOR was used as a query for BLAST (44) by searching in the Protein Data Bank (PDB) and in the RefSeq database. The RefSeq database was selected to limit species redundancy and because it usually contains genomes of non-questionable quality. The searches in the RefSeq database were set up to extract 500 sequences. Sequences with more than 90 % identity were removed using CD-HIT (45, 46), yielding 336 sequences. The sequences extracted from the PDB and from the work from Arndt *et al*. 2019 (11) were added, leading to a total of 369 sequences used for the construction of the tree. The phylogenetic trees were constructed using the maximum likelihood method and were generated with the MEGA program (47) by using an alignment constructed with MUSCLE (48). A total of 200 replicates were used to calculate each node score. The figure was constructed using iTOL (49), and the taxonomy was performed using NCBI taxonomy (50).

The Fig. S6a was constructed using the WebLogo 3 webserver (51), from an alignment performed with Clustal Omega (52) from the 336 sequences obtained by BLAST and CD-HIT.

### Crystallisation and structure determination

Crystallisation was performed anaerobically by initial screening at 20 °C using the sitting drop method on 96-Well MRC 2-Drop polystyrene Crystallisation Plates (SWISSCI) in a Coy tent containing an N_2_/H_2_ (97:3%) atmosphere. The reservoir chamber was filled with 90 μl of crystallisation condition (SG1™ crystallisation screen from Molecular Dimensions), and the crystallisation drop was formed by spotting 0.55 μl of purified protein with 0.55 μl of precipitant. The protein was crystallised at 8.6 mg.ml^-1^ in storage buffer, and the precipitant solution contained 200 mM MgCl_2_ x 6 H_2_O, 100 mM Bis-Tris pH 6.5, 25% (w/v) PEG 3,350. The crystals were soaked in the crystallisation solution supplemented with 25% (v/v) glycerol for a few seconds before freezing in liquid nitrogen.

### Data collection and structural analysis

The diffraction experiments used for the deposited model were performed at 100 K on the beamline PROXIMA-1 from SOLEIL. The initial determination of the structure was performed using diffraction data collected on the beamline P11 from DESY. The data from SOLEIL were preferred for refinement based on a lower pseudo-translational symmetry. Indeed, the analysis of the Patterson function by Xtriage revealed a significant off-origin peak that is 66.05 % of the origin peak, indicating pseudo-translational symmetry. The data were processed and scaled with *autoPROC* (53). The data presented anisotropy (along the following axes: *a* = 1.92 Å, *b* = 1.57 Å, and *c* = 1.71 Å) and were further processed with *STARANISO* correction integrated with the *autoPROC* pipeline (54) (*STARANISO*. Cambridge, United Kingdom: Global Phasing Ltd.).

The structure was solved by processing a single-wavelength anomalous dispersion experiment at the W L3-edge (Fig. S5, Table S1) using the *CRANK-2* program. The model was manually built *via COOT* (55) and refined with *PHENIX* (version 1.20.1-4487). The last refinement steps were performed by refining with a translation libration screw (TLS), and models were validated by the MolProbity server (56). The model was refined with hydrogens in the riding position. Hydrogens were omitted in the final deposited models. The PDB ID code of the structure is 9G7J. Data collection and refinement statistics for the deposited models are listed in Table S1. All figures were generated and rendered with PyMOL (Version 2.2.0, Schrödinger, LLC, New York, NY, United States).

The data presented in Fig. 1d were obtained from an analysis of the PDB. The shortest distance between the pterin and the closest [4Fe-4S] was measured, and an average of the measures obtained in the asymmetric unit was calculated. This average was indicated in the figure (*Ca*AFOR) or used to calculate an average for the group of enzymes. The structures used were 1) *Ca*AFOR (9G7J), 2), 1B25, 1AOR, 8C0Z, 6X1O, 6X6U and 4Z3W (aldehyde oxidases, AOX), 3) 5T5I, 2VPW, 1AA6, 7BKB, 7VW6, 7E5Z, 2E7Z, 5E7O, 4YDD, 2JIO, 2JIM, 3EGW, 3IR7, 1R27, 1Q16 and 3IR5 (bis-PGD enzymes) and 4) 5G5G, 5G5H, 5Y6Q, 1RM6, 4ZOH, 1DGJ, 1T3Q, 4C7Y, 1SIJ, 7OPN, 5EPG, 7DQX, 4UHW, 7PX0, 1N5W, 1ZXI and 8GY3 (Xanthine oxidases).

### High-resolution clear native PAGE (hrCN PAGE) and size-exclusion chromatography

The hrCN PAGE protocol was adapted from Lemaire *et al*. (2018) (57). Glycerol was added to the sample at a final amount of 20 % (v/v). Ponceau S at a final concentration of 0.001 % (w/v) served as a marker to follow the migration. The buffer composition for the electrophoresis cathode was the following: 50 mM Tricine, 15 mM Bis-Tris/HCl pH 7.0, 0.05 % (w/v) sodium deoxycholate, 2 mM DTT, and 0.01 % (w/v) dodecyl maltoside, whilst the anode buffer contained 50 mM Bis-Tris/HCl, PH 7.0 and 2 mM DTT. An 8– 15 % linear polyacrylamide gradient gel was used, and electrophoresis was run under a N_2_/CO_2_ (90:10 %) atmosphere with a constant 40 mA current (PowerPac^™^ Basic Power Supply, Bio-Rad). After electrophoresis, protein bands were visualised with Ready Blue^™^ Protein Gel stain (Sigma Aldrich, Hamburg, Germany). The native protein ladder used is NativeMark^™^ Unstained Protein Standard (Thermo Fischer Scientific, Driesch, Germany).

The determination of the oligomeric state by gel filtration was performed on a Superdex 200 10/300 GL and a Superose 6 Increase 10/300 GL (GE Healthcare, Munich, Germany) in 25 mM Tris/HCl pH 7.6, 10 % (v/v) glycerol and 2 mM DTT at a flow rate of 0.4 ml.min^-1^ in an anaerobic Coy tent containing an N_2_/H_2_ (97:3 %) atmosphere at 20 °C. A high molecular weight range gel filtration calibration kit (GE Healthcare, Munich, Germany) was used as the protein standard.

### Activity measurements

The enzyme preincubation for reactivation was performed in 50 mM Tris/HCl pH 8.0 at 20 °C in an anaerobic chamber filled with an N_2_ (100 %) atmosphere. A *Ca*AFOR final concentration in the preincubation mix ranged between 0.16 and 4.7 µM was used. The used ferredoxin final concentration in the preincubation mix ranged from 0 to 250 µM ferredoxin. When indicated, 500 µM Ti(III) citrate, 500 µM ferricyanide, 500 µM dithionite and 500 µM sulphide final concentration in the preincubation mix were used.

Aldehyde:MV oxidoreduction activity was assayed by measuring the reduction of MV at 600 nm in 50 mM Tris/HCl pH 8.0, 5 mM MV, at 37 °C in a BMG Labtech FLUOstar Omega Microplate reader in an anaerobic chamber filled with an N_2_ (100%) atmosphere. When indicated, BV was used instead of MV at the same concentration. The final protein concentration ranged from 1.6 to 66 nM. An experimentally determined molar extinction coefficient of 17,213 M^−1^.cm^-1^ for reduced MV and 6,384.2 M^−1^.cm^-1^ for reduced BV were used for calculation.

As for other characterised aldehyde oxidases, BV acts as more suitable electron acceptor for aldehyde oxidation than MV (Fig. S10b). As the measured viologen-based activities are used to compare different conditions, the used artificial electron acceptor is relatively meaningless, the important stated activities being measured with the physiological electron shuttle ferredoxin.

All measurements have been done at least in triplicates consisting of three different reaction mixtures using the same reactivated enzyme and are presented in μmol of reduced MV/BV/ferredoxin/NADH per minute per mg of protein (as indicated). The data used for calculation are subtracted from the baseline measured without substrate addition. The kinetic parameters for formaldehyde could not be estimated as the necessary concentrations of the aldehyde triggered enzyme aggregation.

The effect of O_2_ on the preincubated *Ca*AFOR could not be assessed due to the complexity of the experimental setup (e.g., cross-reaction with Ti(III) citrate, ferredoxin and sulphide). The CO sensitivity was assayed using a Cary 60 UV–Vis spectrophotometer (Agilent Technologies) in 600 µl quartz cuvette and at 37 °C. The reactivation was initiated by acetaldehyde addition, and CO was added after the initial measurement of the activity.

Ferredoxin reduction by *Ca*AFOR and CODH/ACS was assayed in quartz cuvette and at 37 °C. The measurement was performed at 50 µM ferredoxin, monitoring the activity at 390 nm, considering a single electron transferred to ferredoxin and with an experimentally determined molar extinction coefficient of 11,095.4 M^-1^.cm^-1^ (obtained by subtraction of the absorbance of aerobically oxidised and dithionite-reduced ferredoxin).

Assays with gasses (100% CO or H_2_, 10% CO) were carried out in a 600 µl quartz cuvette containing a 400 µl reaction mixture. The gas (0.1 ml) was anaerobically injected with a syringe. The data used for calculation are subtracted from the baseline measured before gas addition.

Coupled enzymatic assays were measured in a quartz cuvette and at 37 °C. A concentration of *Ca*AFOR and CODH/ACS of 32 and 1,214 nM were used, respectively. A final concentration of ferredoxin ranging from 2 to 10 µM was used. The ethanol dehydrogenase from *S. cerevisiae* (15.6 U.ml^-1^ final) was used to reduce acetaldehyde, 0.2 mM NADH being used as an electron donor. Oxidation of NADH was used to monitor activity, followed at 340 nm, calculated considering a mole of oxidised NADH per mole of reduced acetaldehyde and using an experimentally determined molar extinction coefficient of 5,297.9 M^-1^.cm^-1^ for calculation. The addition of 0.1 ml CO (100% or 10%) was used to initiate the reaction. The data used for calculation are subtracted from the baseline measured before CO addition.

## Author contributions

Organism cultivation, enzyme purification, biochemical characterisation, activity assays and crystallisation were performed by O.N.L. and M.B. Peptide identification by mass spectrometry was performed by A.S. O.N.L. performed phylogenetic analyses. X-ray data collection was performed by O.N.L. and T.W. Data processing, model building, structure refinement, validation and deposition were performed by O.N.L. and T.W. Structures were analysed by O.N.L. and T.W. The paper was written by O.N.L. and T.W. with contributions and final approval of all co-authors.

## Acknowledgements

We thank the Max Planck Institute for Marine Microbiology and the Max Planck Society for their continuous support. The work was additionally supported by the Deutsche Forschungsgemeinschaft priority program 1927, “Iron-Sulfur for Life” WA 4053/1-1. We also thank Christina Probian and Ramona Appel for their continuous support in the Microbial Metabolism laboratory. We thank the DESY and SOLEIL Synchrotrons, especially the staff of beamline PROXIMA-1 from SOLEIL and P11 at DESY.

