## Supplementary information for "Carbon monoxide-driven bioethanol production operates via a tungsten-dependent catalyst"

### Supplementary data

This file contains:

Supplementary figures Fig. S1-14

Supplementary table 1

References

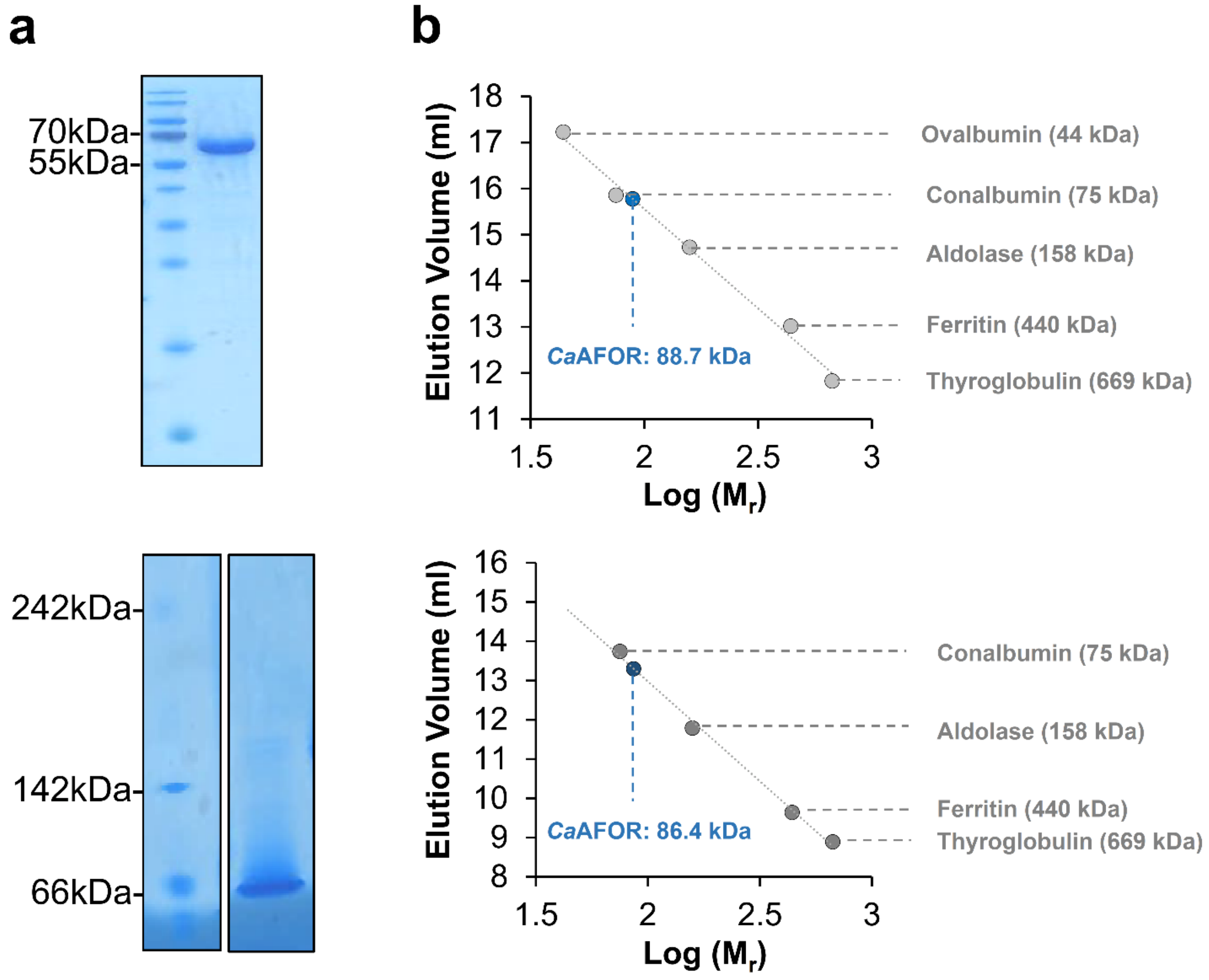


**Fig. S1. Size determination of the AFOR from *C. autoethanogenum*.** **a**, Denaturing (top) and native (bottom) electrophoresis profile of the purified *Ca*AFOR (3 µg). **b**, Determination of molecular weight by size exclusion chromatography. The molecular weight of *Ca*AFOR (blue circles) was determined using a Superose 6 increase 10/300 GL (top), a Superdex 200 10/300 increase GL (bottom) and a protein standard calibration kit (grey circles). *Ca*AFOR has an expected molecular weight of 66,335 Da based on its sequence.

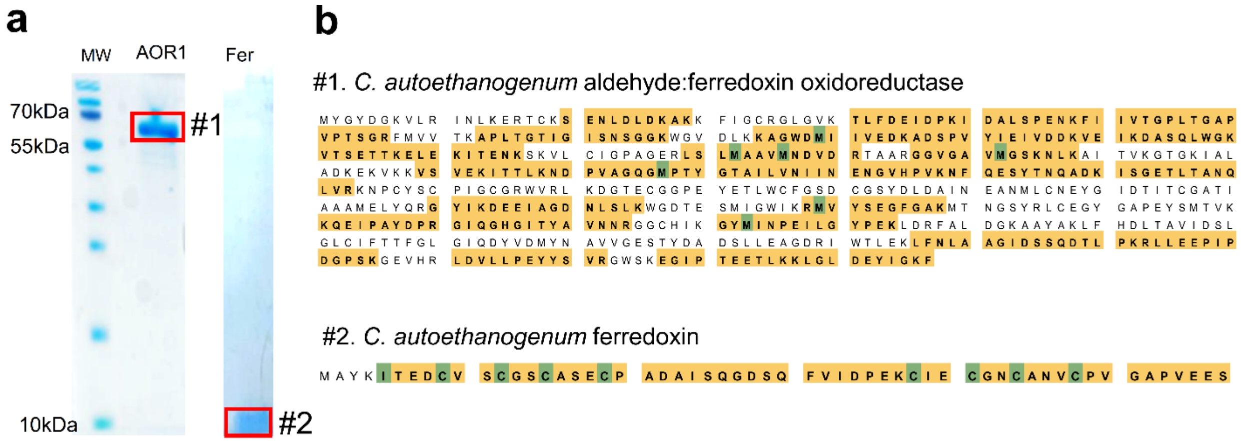


**c**

| Sample | Protein name | Gene name | Acc. No. | Molecular weight | Mass Spectrometry Analysis matched | | |
|---|---|---|---|---|---|---|---|
| | | | | | Number of unique peptides | Number of spectra | Sequence coverage |
| #1 | Aldehyde:ferredoxin oxidoreductase* | *Aor1* | WP_013238665 | 66 kDa | 37 | 225 | 55% |
| #2 | Ferredoxin | *fer* | WP_013236834/ OVY50687.1 | 6 kDa | 2 | 39 | 93% |

**Fig. S2. Mass Spectrometric identification of AOR1 (*Ca*AFOR) and ferredoxin proteins purified from *C. autoethanogenum*.** **a**. Coomassie-stained SDS gel images of purified *Ca*AFOR and ferredoxin purifications. Gel regions analysed by mass spectrometry are designated with red squares. Selected MS markers are shown on the left-hand side. **b**. Distribution of peptides detected by mass spectrometry for *Ca*AFOR and ferredoxin in samples #1 and #2. MS-matched peptides are highlighted in yellow, and amino acids detected carrying modifications (oxidation, carbamidomethylation, acetylation) are marked in green. **c**. MS-identification details for both proteins in the corresponding gel regions (a). * Only traces of the second AFOR isoform or *C. autoethanogenum* WP_013238675.1 (two peptides) were detected in the band #1.

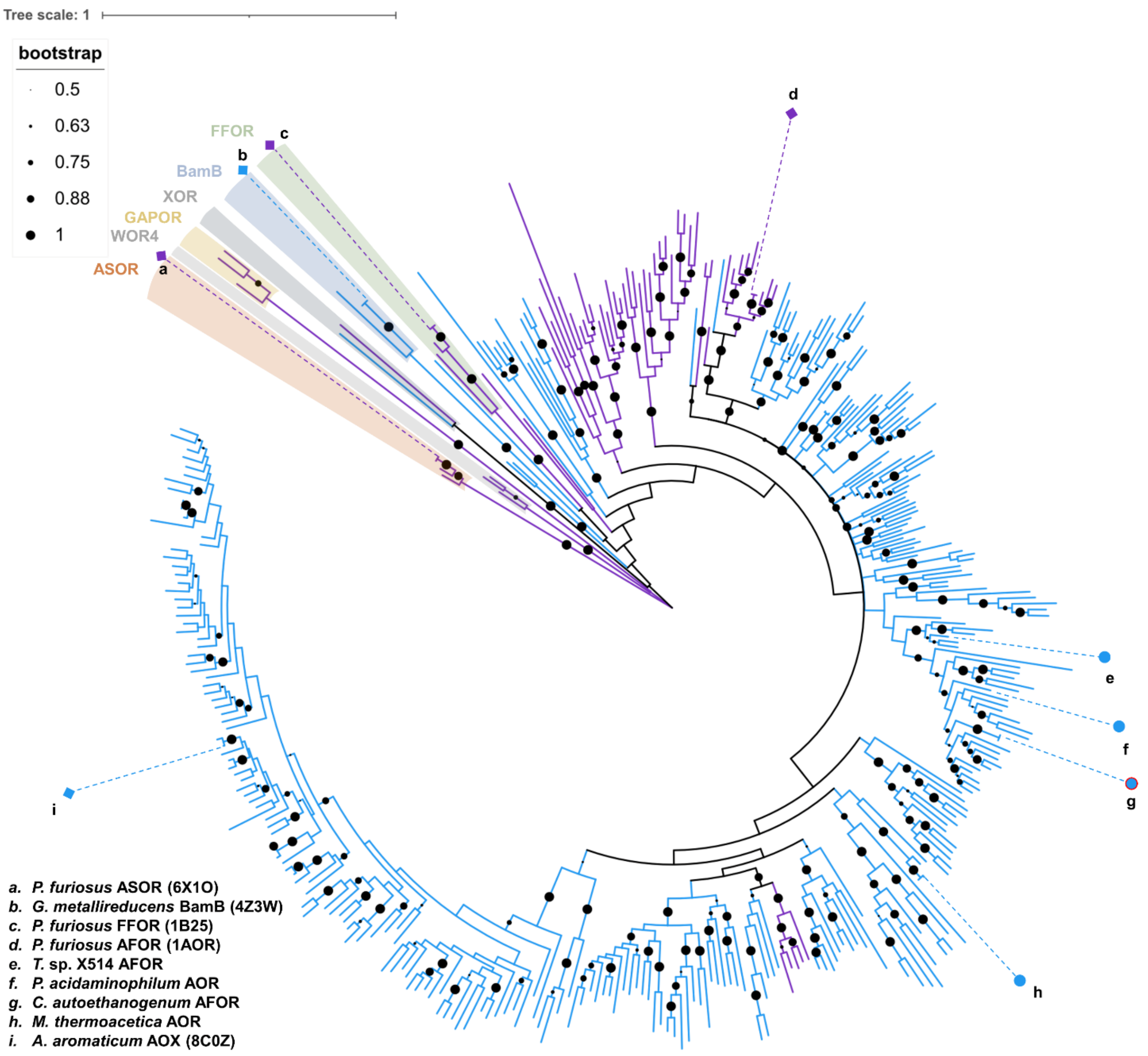


**Fig. S3. Phylogenetic tree of aldehyde oxidases.** The maximum likelihood phylogenetic tree was built by using 369 sequences composed of (*i*) *Ca*AFOR homologues presenting 90 % or less identity, (*ii*) sequences of the structurally characterised homologues, and (*iii*) sequences used in the phylogenetic analysis from Arndt *et al.* 2019 (1). Node scores were calculated on 200 replicates. Branches are coloured blue (bacteria) or purple (archaea), and enzyme types are indicated (1). The closest branch of the characterised enzymes is indicated and labelled. Structurally characterised enzymes are indicated as squares, and biochemically characterised enzymes are coloured in circles depending on the taxonomy.

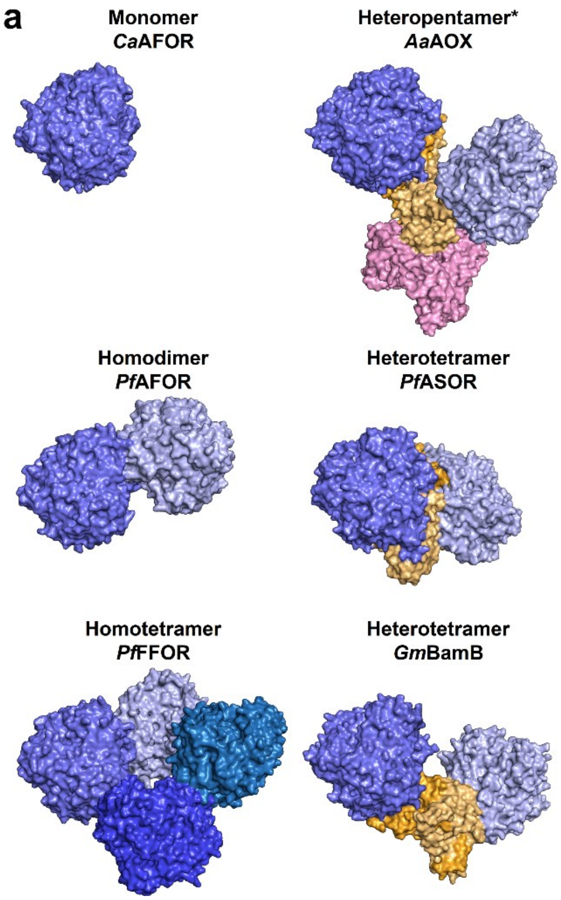


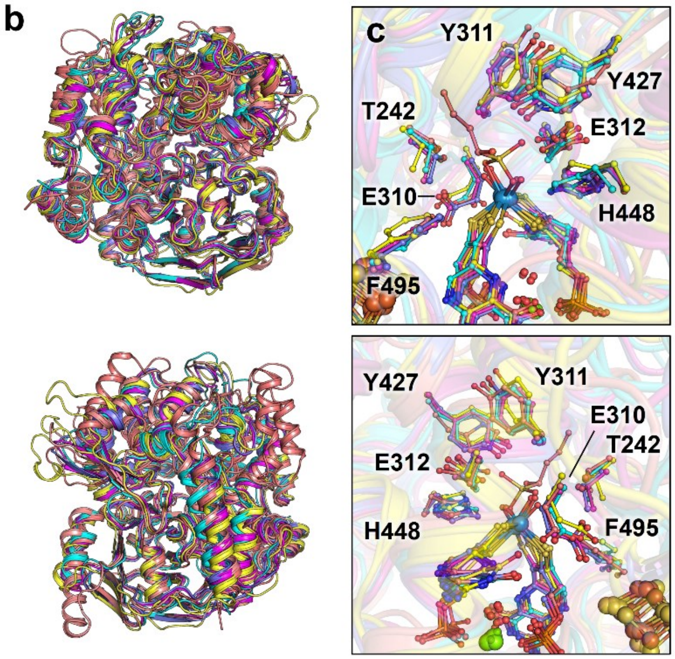


**d**

| Structure | r.m.s.d. (Cα) |
|---|---|
| ***Ca*AFOR, 9G7J** | 0.000 (609) |
| ***Pf*AFOR, 1AOR** | 0.516 (510) |
| ***Pf*FFOR, 1B25** | 1.221 (461) |
| ***Aa*AOX, 8C0Z** | 1.294 (559) |
| ***Pf*ASOR, 6X1O** | 1.377 (435) |
| ***Gm*BamB, 4Z3W** | 1.380 (381) |

**Fig. S4. Structural organisation of aldehyde oxidases-containing complexes. a**, Organisation of the structurally characterised aldehyde oxidases. Proteins are shown as surfaces with aldehyde oxidases, electron transfer subunits and $NAD^+$-reducing subunits in shades of blue, orange and pink, respectively. The asterisk for the heteropentameric *Aa*AOX highlights the property of this complex to generate nanowire-like structures. **b**, **c**, Superposition of the aldehyde oxidases subunits. **b, c**, Proteins are shown as cartoon coloured as follows: *Ca*AFOR (9G7J, slate), the AFOR from *P. furiosus* (1AOR, pink), the FFOR from *P. furiosus* (1B25, cyan), the AOX from *A. aromaticum* (8C0Z, yellow) and the ASOR from *P. furiosus* (6X1O, salmon). Alignment root mean square deviation (r.m.s.d.) and aligned Cα are given in **d**. The numbering is from *Ca*AFOR. **c**, The tungstopterin cofactor and neighbouring residues are shown as balls and sticks with oxygen, nitrogen, sulphur, phosphorus, magnesium, iron and tungsten coloured red, blue, light yellow, light orange, green, orange and grey blue, respectively. **d**, R.m.s.d. of the structurally characterised aldehyde oxidase subunits.

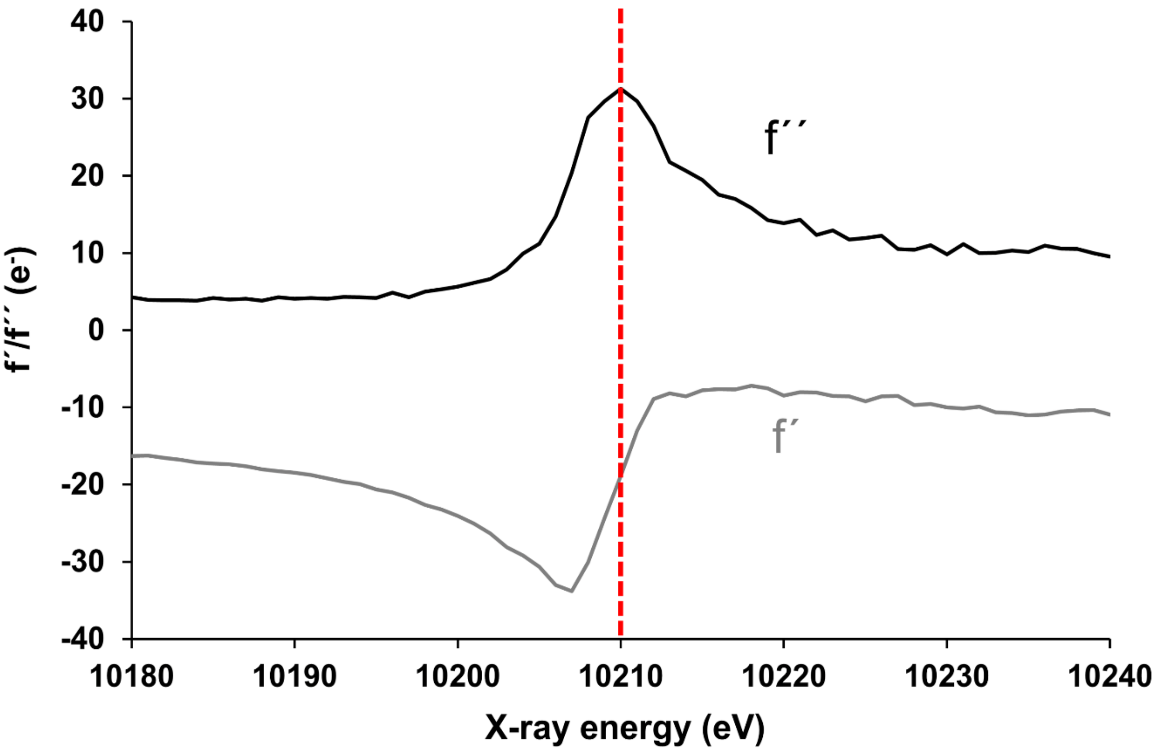


**Fig. S5. Fluorescence scan from a *Ca*AFOR protein crystal.** The fluorescence counts are given as a function of energy (eV). The dashed red line indicates the energy used for data collection (10,210 eV).

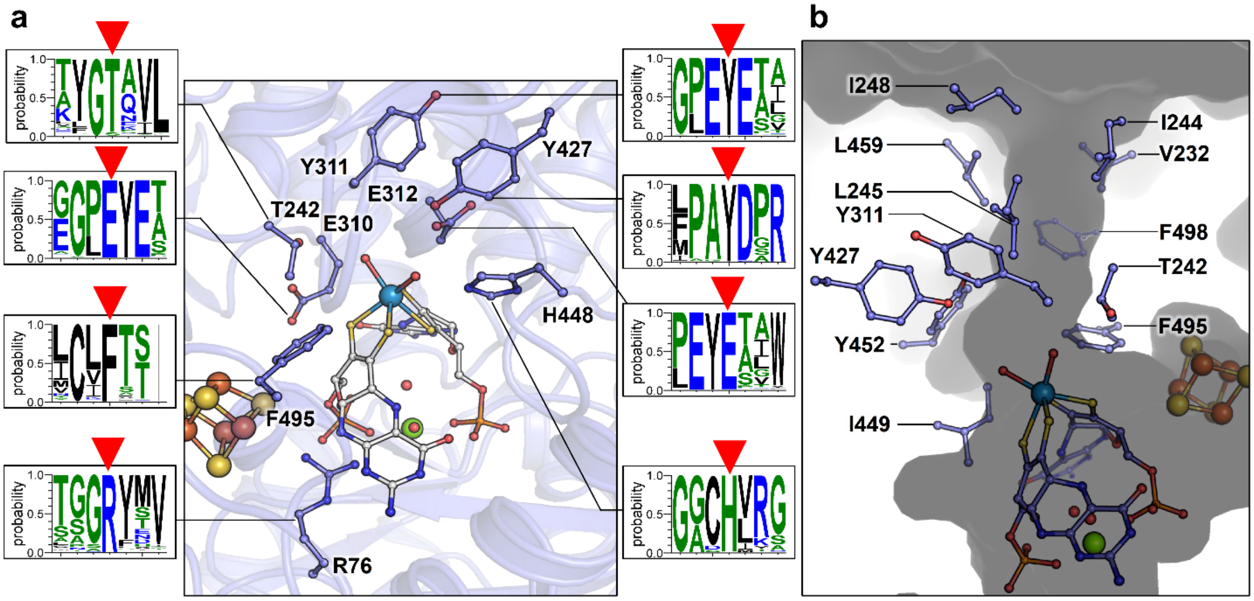


**Fig. S6. Residue conservation in the vicinity of the catalytic tungsten atom.** The structure of *Ca*AFOR is shown as cartoon with cofactors and neighbouring residues shown as balls and sticks. Schematic representation of residue conservation (constructed with Weblogo 3 (2)) using the 336 sequences extracted from the RefSeq database (Fig. S3) are shown in the side panels. A red arrow indicates the residue of interest. **b**, Access to the active site. The protein is shown as a grey surface. The residues coordinating the tunnels leading to the active sites are shown as balls and sticks. **a** and **b**, Oxygen, nitrogen, sulphur, phosphorus, magnesium, iron, and tungsten are coloured red, blue, light yellow, light orange, green, orange and grey blue, respectively.

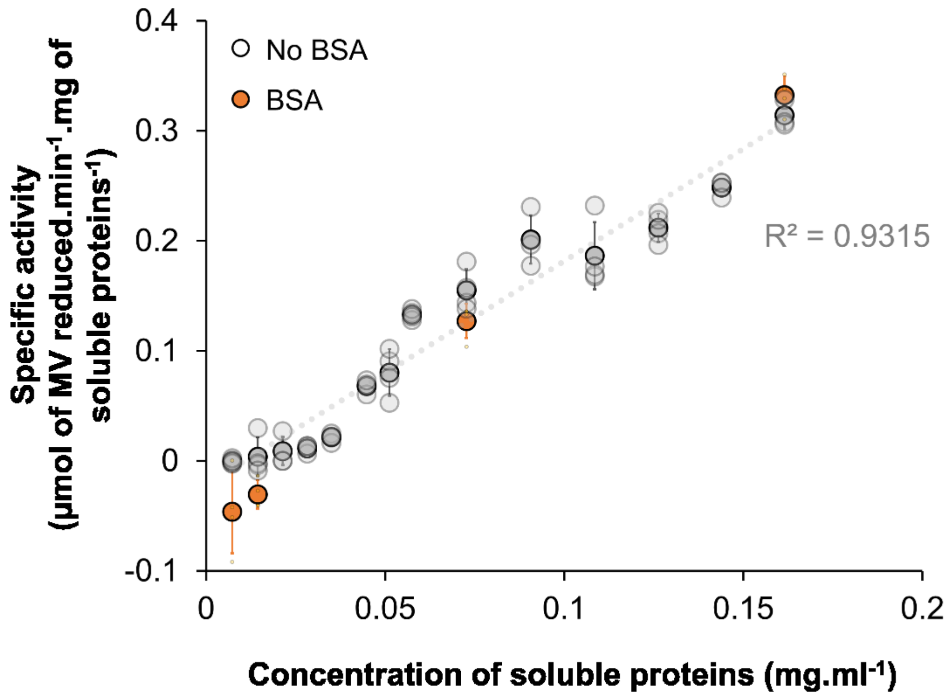


**Fig. S7. AFOR activity in soluble extracts depending on the extract concentration.** The activity was measured in the presence (orange circles) or the absence (white circles) of Bovine Serum Albumin. Average and standard deviation are shown, with individual data shown as transparent grey dots. MV is used as an electron acceptor.

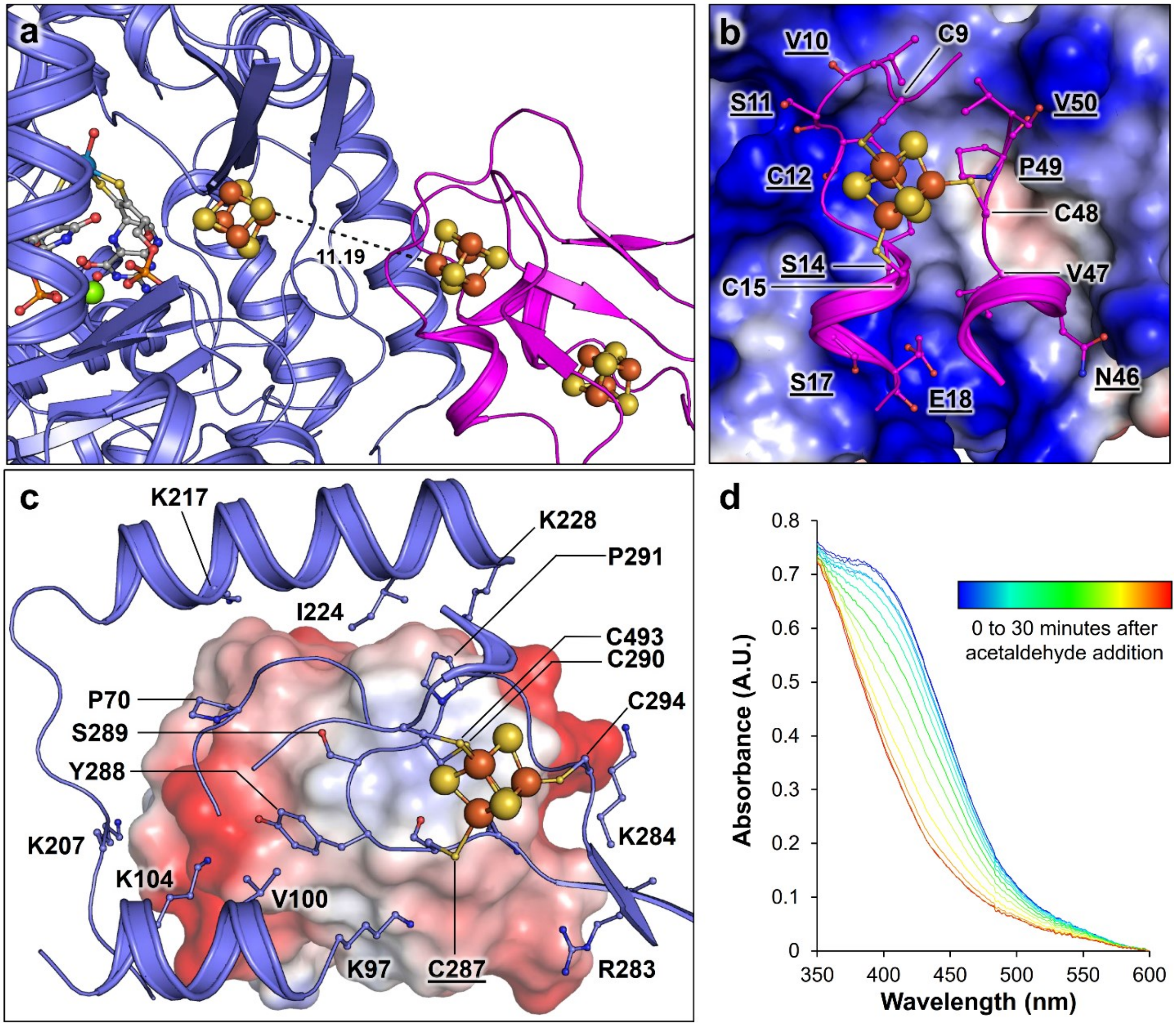


**Fig. S8. Ferredoxin docking and reduction by *Ca*AFOR. a**, Distances (indicated in Å) between ferredoxin and *Ca*AFOR clusters in the AlphaFold2 model presented in Fig. 3b. **b, c**, Mapping of the *Ca*AFOR-ferredoxin interface. The ferredoxin docking on *Ca*AFOR (shown as surface coloured by electrostatic charges) and the *Ca*AFOR docking on ferredoxin in a different orientation (shown as surface coloured by electrostatic charges) are shown in **b** and **c**, respectively. **a – c**, Proteins are shown as cartoons, and *Ca*AFOR and ferredoxin are coloured slate and pink, respectively. Residues involved in cofactor coordination or docking stabilisation are shown as balls and sticks. Residues interacting with main chains are underlined. Cofactors are shown as balls and sticks, with carbon, oxygen, nitrogen, sulphur, phosphorus, magnesium, iron and tungsten coloured white (as the respective proteins for residues), red, blue, light yellow, light orange, green, orange and grey blue, respectively. **d**, Ferredoxin reduction by *Ca*AFOR, monitored by measuring the absorbance spectrum every 2.5 minutes. Spectra are coloured blue to red from the first to the last time point.

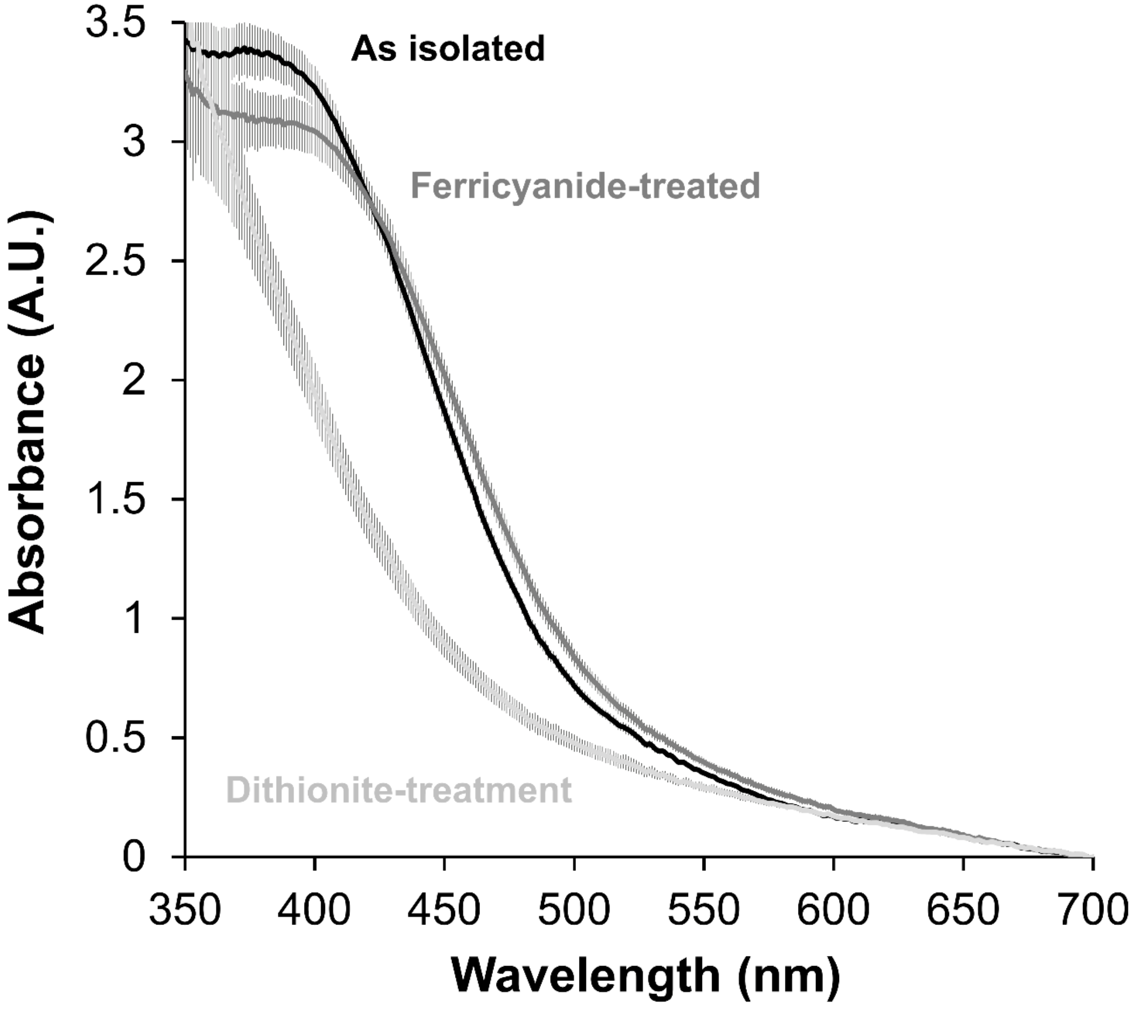


**Fig. S9. Redox state of the isolated ferredoxin.** The absorbance spectrum of the ferredoxin as isolated or treated with an oxidising (ferricyanide) or reducing (dithionite) agent. Average and standard deviation of three distinct spectra are displayed.

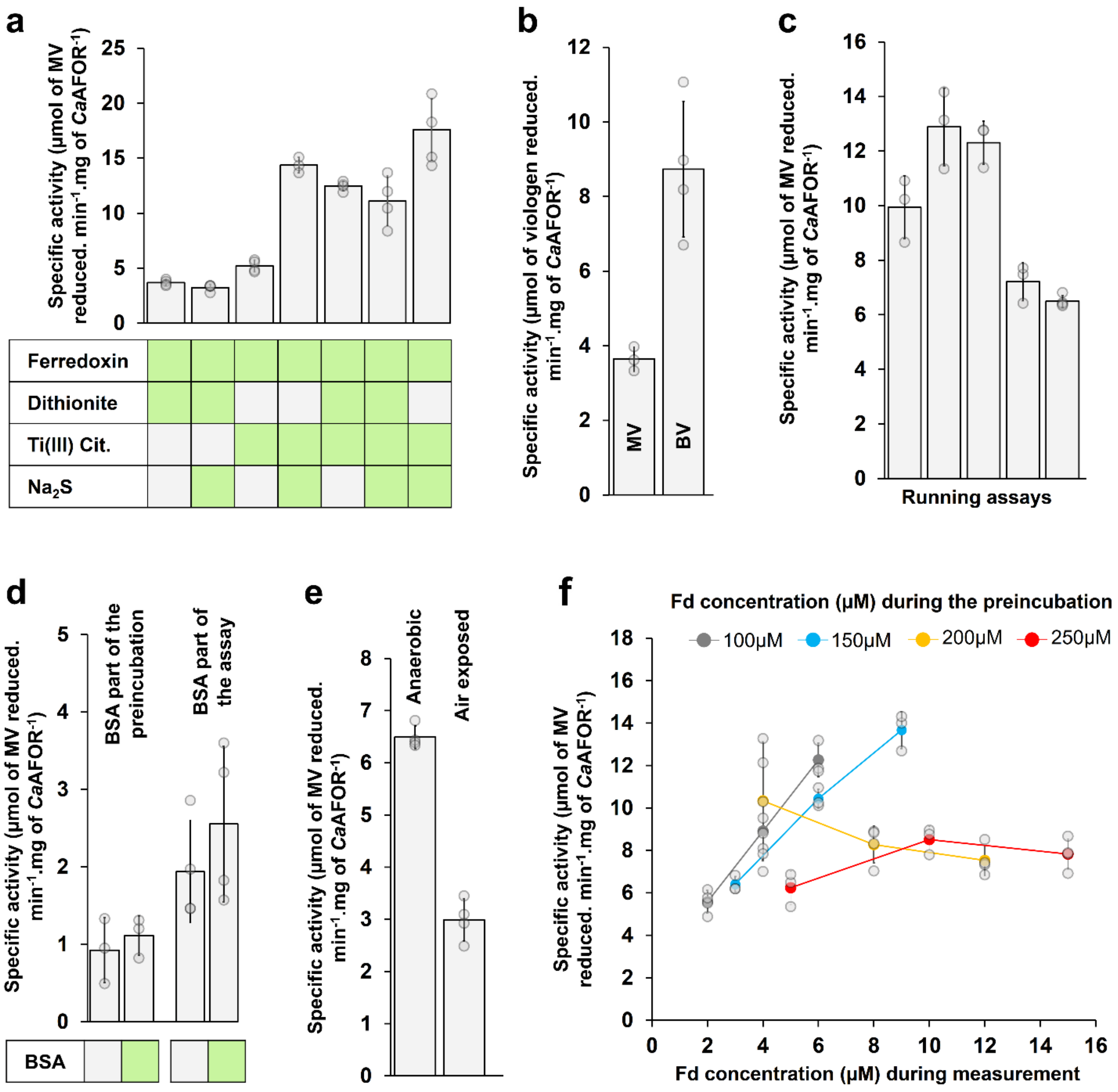


**Fig. S10. Enzyme reactivation and activity.** **a**, Reactivation of *Ca*AFOR with 250 µM ferredoxin, 500 µM dithionite, 500 µM Ti(III) citrate and/or 500 µM sulphide. **b**, Activity of *Ca*AFOR with MV and BV (both at 5 mM) as electron acceptor. **c**, Activity measurements in multiple reactivations of the same pool of frozen enzyme. **d**, Effect of BSA during reactivation (left) or activity measurements (right). **e**, Activity measured after overnight incubation under anaerobic or aerobic conditions. Reactivation procedure and activity measurements were performed in anaerobic conditions for both. **f**, Impact of the ferredoxin concentration during reactivation and activity measurements. The enzyme was reactivated with different concentrations of ferredoxin (represented by different colours). By dilution, three different final concentrations of ferredoxin were assessed during activity measurements for each reactivation procedure. **a** and **c-f**, MV serves as electron acceptor. **a-f**, Average and standard deviation are shown, with individual data shown as transparent grey dots.

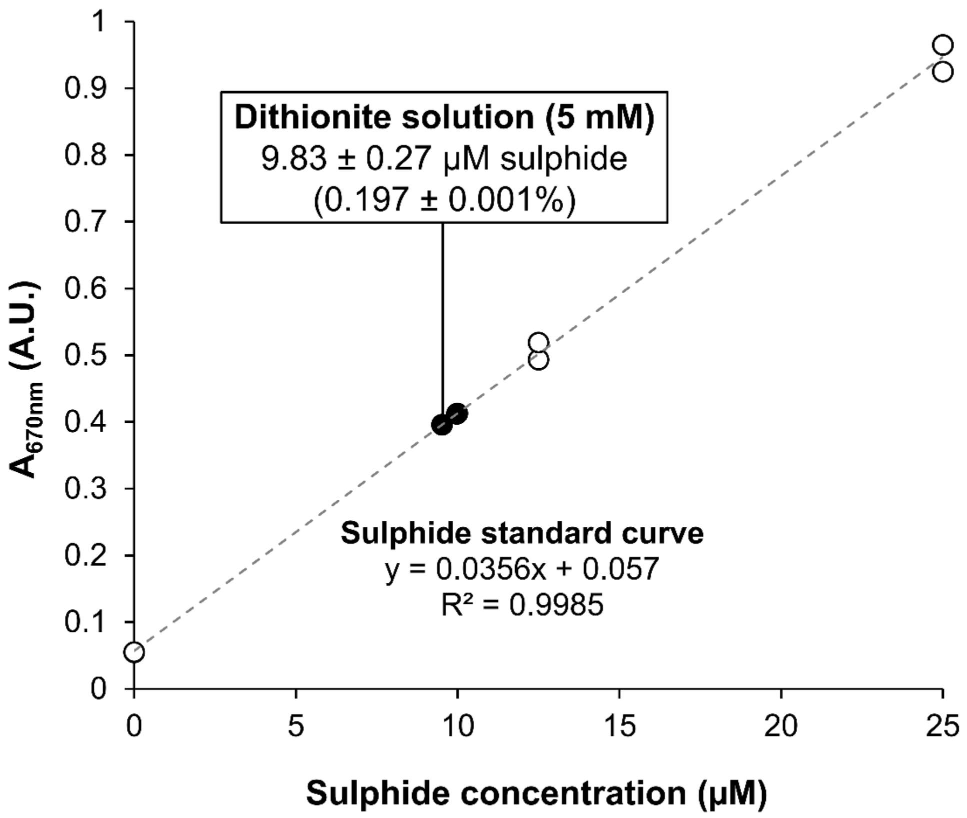


**Fig. S11. Sulphide quantification in the dithionite solution used in the assay (Fig. S10a).** Absorbance at 670 nm is plotted. Measures of the sulphide standards and dithionite samples are shown as white and black circles, respectively.

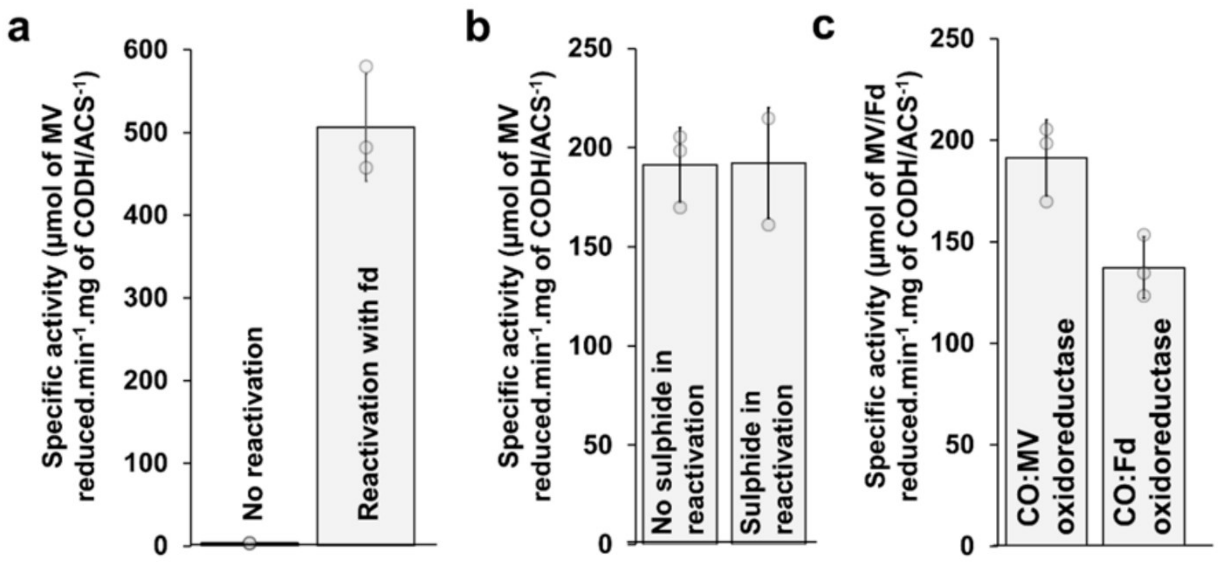


**Fig. S12. Reactivation and activity of the CODH/ACS. a**, Specific activity of the CODH/ACS(3) without and with reactivation by incubation with ferredoxin and Ti(III) citrate. **b**, Impact of sulphide on the CODH/ACS reactivation process. **c**, CO:MV and CO:ferredoxin oxidoreductase activity of the CODH/ACS complex(3) after reactivation. **a-c**, Average and standard deviation are shown, with individual data shown as transparent grey dots.

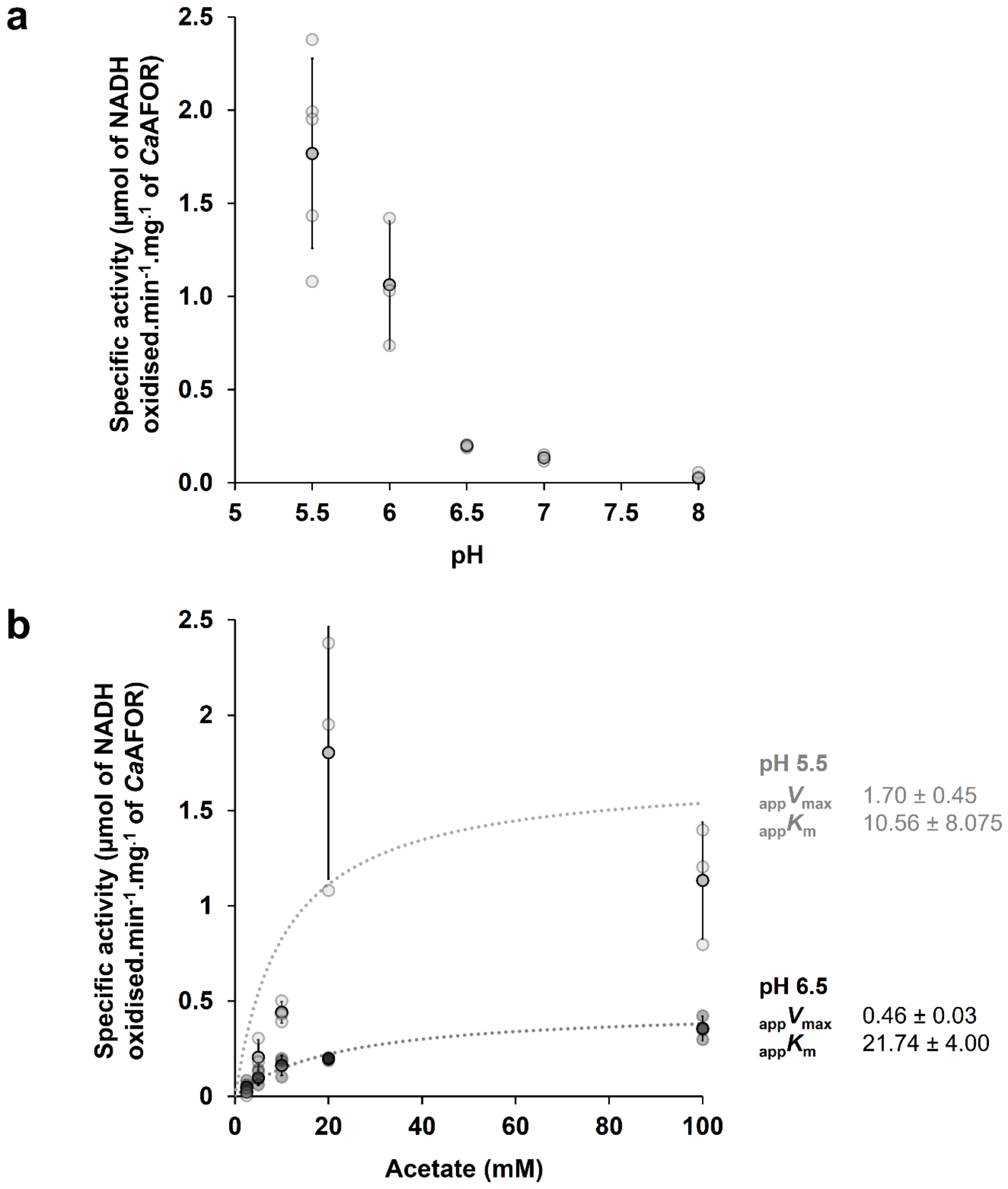


**Fig. S13. pH and concentration impact on the monitored coupled assay. a**, **b**, Specific activity (per mg of *Ca*AFOR) in the coupled assay depending on pH (**a**) and sodium acetate concentration (**b**). b, Measurements were performed at pH 5.5 (light grey) or 6.5 (dark grey). With a pKa of 4.76 for acetic acid/acetate, the estimated ${}_{app}K_m$ of *Ca*AFOR for acetic acid would represent around 1.63 and 1.18 mM in the solution at pH 5.5 and 6.5, respectively. **a, b**, Average and standard deviation are shown, with individual data shown as transparent grey dots.

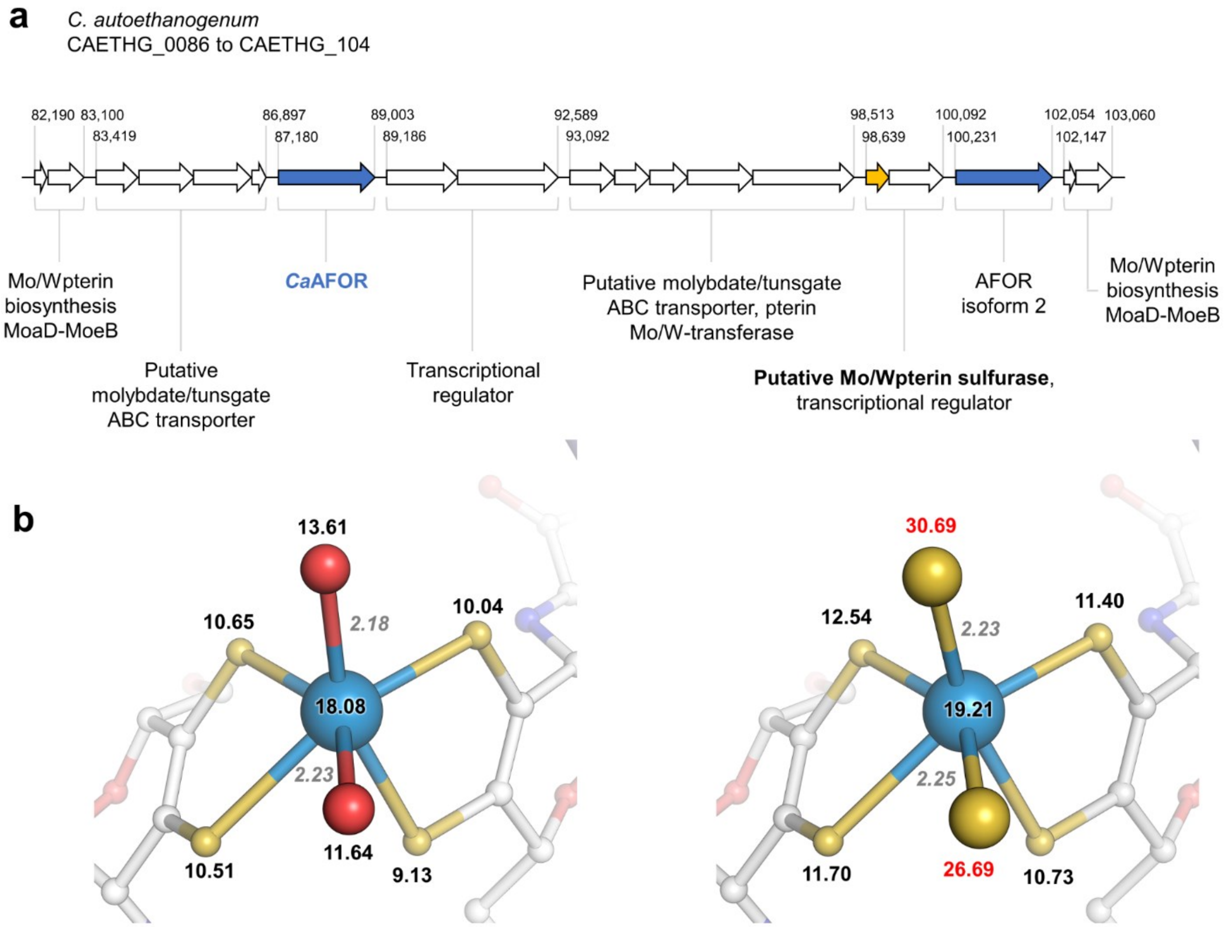


**Fig. S14. Genomic environment and sulphur ligand in *Ca*AFOR. a**, Genomic environment of the genes coding for the *Ca*AFOR and the second AFOR isoform of *C. autoethanogenum* (slate). Genes are shown as arrows with length depending on the gene length. Adjoining arrows represent operonic structures predicted by the Operon Mapper webserver (4). The gap between predicted operons is not representative of the actual sequence length. The first and last positions of the predicted operons are indicated, such as the predicted function of the encoded proteins. The putative W/Mo-pterin cofactor sulfurase is coloured orange. **b**, Distances and b-factors of the W atom and coordinating ligands when S atoms substitute the O ligands. The cofactor and ligands are shown as balls and sticks with carbon, oxygen, nitrogen, sulphur and tungsten coloured white, red, blue, light yellow and grey-blue, respectively. B-factors are coloured black and distances in grey (given in Å). Modelled S atoms have a b-factor that is inconsistent with the atoms from the environment. As reactivation of the enzyme can be performed without sulphur sources, the putative S-ligand atom in the active site could not be necessary for activity, similar to the sulfido ligand of the Mo-dependent trimethylamine-N-oxide reductase (5). The absence of this ligand sensitive to oxidation in some enzymes may explain the relative tolerance to the oxygen of aldehyde oxidases of facultative aerobic bacteria (1). The higher electronegativity and charge density of sulphur compared to oxygen could be more favourable in reaction, explaining why sulphide enhances enzyme activity without being necessary.

**Table S1. X-ray analysis statistics for the structure of *Ca*AFOR.**

| **Data collection** | |
|---|---|
| Synchrotron source | SOLEIL, PROXIMA-1 |
| Wavelength (Å) | 1.21434 |
| Space group | $P2_122_1$ |
| Resolution (Å) | 100.44 − 1.59 (1.74 −1.59) |
| Cell dimensions | |
| a, b, c (Å) | 64.59, 100.44, 176.59 |
| α, β, γ (°) | 90, 90, 90 |
| $R_{merge}$(%)[a] | 17.3 (172.5) |
| $R_{pim}$ (%)[a] | 7.2 (74.6) |
| $CC_{1/2}$ [a] | 0.997 (0.641) |
| $I/\sigma_I$ [a] | 8.4 (1.6) |
| Spherical completeness[a] | 75.7 (17.1) |
| Ellipsoidal completeness[a] | 95.1 (63.1) |
| Redundancy[a] | 12.7 (12.2) |
| Nr. unique reflections[a] | 116,033 (5,802) |
| **Refinement** | |
| Resolution (Å) | 50.79 − 1.59 |
| Number of reflections | 115,999 |
| $R_{work}/R_{free}$[b] (%) | 16.16/17.79 |
| Number of atoms | |
| Protein | 9,378 |
| Solvent and ligands | 128 |
| Water | 1504 |
| Mean B-value (Å$^2$) | 19.29 |
| Molprobity clash score, all atoms | 1.37 |
| Ramachandran plot | |
| Favoured regions (%) | 95.45 |
| Outlier regions (%) | 0.17 |
| r.m.s.d.[c] bond lengths (Å) | 0.011 |
| r.m.s.d.[c] bond angles (°) | 1.467 |
| **PDB ID code** | 9G7J |

[a] Values relative to the highest resolution shell are within parentheses. [b] $R_{free}$ was calculated as the $R_{work}$ for 5% of the reflections that were not included in the refinement. [c] r.m.s.d., root mean square deviation.

### References.
